# Fine-Scale Adaptations to Environmental Variation and Growth Strategies Drive Phyllosphere *Methylobacterium* Diversity

**DOI:** 10.1101/2021.06.04.447128

**Authors:** Jean-Baptiste Leducq, Émilie Seyer-Lamontagne, Domitille Condrain-Morel, Geneviève Bourret, David Sneddon, James A. Foster, Christopher J. Marx, Jack M. Sullivan, B. Jesse Shapiro, Steven W. Kembel

## Abstract

*Methylobacterium* is a prevalent bacterial genus of the phyllosphere. Despite its ubiquity, little is known about the extent to which its diversity reflects neutral processes like migration and drift, versus environmental filtering of life history strategies and adaptations. In two temperate forests, we investigated how phylogenetic diversity within *Methylobacterium* was structured by biogeography, seasonality, and growth strategies. Using deep, culture-independent barcoded marker gene sequencing coupled with culture-based approaches, we uncovered a considerable diversity of *Methylobacterium* in the phyllosphere. We cultured different subsets of *Methylobacterium* lineages depending upon the temperature of isolation and growth (20 °C or 30 °C), suggesting long-term adaptation to temperature. To a lesser extent than temperature adaptation, *Methylobacterium* diversity was also structured across large (>100km; between forests) and small geographical scales (<1.2km within forests), among host tree species, and was dynamic over seasons. By measuring growth of 79 isolates at different temperature treatments, we observed contrasting growth performances, with strong lineage- and season-dependent variations in growth strategies. Finally, we documented a progressive replacement of lineages with a high-yield growth strategy typical of cooperative, structured communities, in favor of those characterized by rapid growth, resulting in convergence and homogenization of community structure at the end of the growing season. Together our results show how *Methylobacterium* is phylogenetically structured into lineages with distinct growth strategies, which helps explain their differential abundance across regions, host tree species, and time. This works paves the way for further investigation of adaptive strategies and traits within a ubiquitous phyllosphere genus.

**Importance:** *Methylobacterium* is a bacterial group tied to plants. Despite its ubiquity and importance to their hosts, little is known about the processes driving *Methylobacterium* community dynamics. By combining traditional culture-dependent and independent (metabarcoding) approaches, we monitored *Methylobacterium* diversity in two temperate forests over a growing season. On the surface of tree leaves, we discovered remarkably diverse and dynamic *Methylobacterium* communities over short temporal (from June to October) and spatial scales (within 1.2 km). Because we cultured different subsets of *Methylobacterium* diversity depending on the temperature of incubation, we suspected that these dynamics partly reflected climatic adaptation. By culturing strains in lab conditions mimicking seasonal variations, we found that diversity and environmental variations were indeed good predictors of *Methylobacterium* growth performances. Our findings suggest that *Methylobacterium* community dynamics at the surface of tree leaves results from the succession of strains with contrasted growth strategies in response to environmental variations.

## Introduction

The phyllosphere, the aerial parts of plants including leaves, is a microbial habitat estimated to be as vast as twice the surface of the earth (1). Although exposed to harsh conditions including UV radiation, temperature variation, and poor nutrient availability, the phyllosphere harbors a diverse community of microorganisms, of which bacteria are the most abundant (1). A key challenge in microbial ecology and evolution is understanding the evolutionary and ecological processes that maintain diversity in habitats such as the phyllosphere. Bacteria living in the phyllosphere carry out key functions including nitrogen fixation, growth stimulation and protection against pathogens (1–3). At broad spatial and temporal scales, bacterial diversity in the phyllosphere varies as a function of geography and host plant species, potentially due to restricted migration and local adaptation to the biotic and abiotic environment (4–6), leading to patterns of cophylogenetic evolutionary association between phyllosphere bacteria and their host plants (7). Whether those eco-evolutionary processes are important at the scale of several days to several years, as microbes and their host plants migrate and adapt to changing climates, is still an open question (8). Another challenge is to link seasonal variation with plant-associated microbial community dynamics, as shifts in microbial community composition are tighly linked with host plant carbon cycling (9) and ecosystem functions including nitrogen fixation (10). More generally, we understand very little about how the ecological strategies of phyllosphere bacteria vary among lineages and in response to variation in environmental conditions throughout the growing season (9, 11).

Phenotypic traits are often phylogenetically conserved in microbes (12), and these traits influence the assembly of ecological communities through their mediation of organismal interactions with the abiotic and biotic environment (13). Recent work has shown that many microbial traits exhibit phylogenetic signal, with closely related lineages possessing more similar traits, although the phylogenetic depth at which this signal is evident differs among traits (14). Most comparative studies of microbial trait evolution have focused on broad patterns across major phyla and classes (14), although some studies have found evidence for complex patterns of biotic and abiotic niche preferences evolving within genus-level phylogenies (15, 16). Furthermore, to date the majority of studies of the diversity of plant-associated microbes have been based on the use of universal marker genes such as the bacterial *16S rRNA* gene, providing a global picture of long-term bacterial adaptation to different biomes and host plants at broad phylogenetic scales (17). However, these studies lack sufficient resolution to assess the evolutionary processes at finer spatial and temporal scales that lead to the origin of adaptations within microbial genera and species (18, 19).

The Rhizobiales genus *Methylobacterium* (*Alphaproteobacteria*, *Rhizobiales*, *Methylobacteriaceae*) is one of the most prevalent bacterial genera of the phyllosphere, present on nearly every plant (20–22). Characterized by pink colonies due to carotenoid production, *Methylobacterium* are facultative methylotrophs, able to use one-carbon compounds, such as methanol excreted by plants, as sole carbon sources (23, 24). Experimental studies have shown the important roles of *Methylobacterium* in plant physiology, including growth stimulation through hormone secretion (25–27), heavy metal sequestration (27), anti-phytopathogenic compound secretion, and nitrogen fixation in plant nodules (28), sparking increasing interest in the use of *Methylobacterium* in plant biotechnology applications (27, 29, 30). Although up to 64 *Methylobacterium* species have been described (31–39), genomic and phenotypic information was until recently limited to a small number of model species: *M. extorquens, M. populi*, *M. nodulans*, *M. aquaticum* and *M radiotolerans*, mostly isolated from anthropogenic environments, and only rarely from plants (40–44). Aditionaly, *Methylobacterium* was mostly isolated assuming its optimal growth was in the range 25-30 °C (45), an approach that could bias strain collections toward mesophylic isolates to the exclusion of isolates from temperate forests where temperatures typically range from 10 to 20 °C during the growing season (46). Newly available genomic and metagenomic data now allow a better understanding of the distribution of *Methylobacterium* diversity across biomes (31) and suggest that they represent a stable and diverse fraction of the phyllosphere microbiota (22). However, we still understand relatively little about the drivers of the evolution and adaptation of *Methylobacterium* in natural habitats.

In this study, we assessed the diversity of *Methylobacterium* in temperate forests and asked whether *Methylobacterium* associated with tree leaves act as a single unstructured population, or if their diversity is structured by regional factors (*e.g*. a combination of isolation by distance and regional environmental variation) or by niche adaptation (*e.g*. host tree or temperature adaptation) (12). First, we assessed *Methylobacterium* diversity by combining culturing and metabarcoding approaches along with phylogenetic analysis and quantified how this diversity varied across space, time, and environment in the phyllosphere. Second, we quantified the extent of phylogenetic niche differentiation within the genus, with a focus on quantifying the evidence for adaptation to local environmental variation at different spatial, temporal and phylogenetic scales. We hypothesized that distinct phylogenetic lineages would be associated with distinct environmental niches. Third, we quantified *Methylobacterium* growth performance under fine-scale environmental variations, with a focus on temperature, to determine whether fine-scale changes in diversity over space and time might result from environmental filtering of isolates with contrasting growth strategies under local environmental conditions. We found that *Methylobacterium* phyllosphere diversity consisted of deeply branching phylogenetic lineages associated with distinct growth phenotypes, isolation temperatures, and large-scale spatial effects (forest of origin), while finer-scale spatial effects, host tree species, and time of sampling were more weakly and shallowly phylogenetically structured. Over the course of a year, from spring to fall, we observed a homogenization of *Methylobacterium* community structure coinciding with the progressive replacement of isolates with high yield strategy by isolates with rapid growth. Together our results show that this ubiquitous phyllosphere genus is structured into lineages with distinct growth strategies, which helps explain their differential abundance across space and time.

## Methods

### Phylogenetics of plant-associated Methylobacterium diversity

We evaluated the known *Methylobacterium* diversity and its distribution across biomes, with a special emphasis on the phyllosphere. First, we constructed a phylogeny of *Methylobacteriaceae* from the complete nucleotide sequence of *rpoB*, a highly polymorphic housekeeping gene commonly used to reconstruct robust phylogenies in bacteria, because unlikely to experience horizontal gene transfer or copy number variation (47, 48). We retrieved *rpoB* sequences from genomes publicly available in September 2020, including 153 *Methylobacteria*, 30 *Microvirga* and 2 *Enterovirga* (**Dataset S1a**), performed alignment and inferred a consensus phylogeny with MrBayes v. 3.2.7a ((49); **Supplementary Materials and Methods S1)**. For each *Methylobacterium* reference genome, we retrieved the species name and the sampling origin, when available. Additionally, we assigned each genome to a group (A, B, C) according to previously proposed subdivisions (31). We subdivided group A in nine clades (A1-A9; **Supplementary Materials and Methods S1**).

### Study sites and sample collection

The two study forests were located at the Gault Nature Reserve (Mont Saint-Hilaire, Quebec, Canada; 45.54 N 73.16 W), here referred as MSH, an old forest occupying Mount Saint-Hilaire, and the *Station Biologique des Laurentides* (Saint-Hippolyte, Quebec, Canada; 45.99 N 73.99 W), here referred to as SBL, a mosaic of natural wetlands, xeric and mesic forests (**Figure 1**, **Dataset S1b**). In August 2017, for the purpose of a pilot survey, we collected leaves from the subcanopy (3-5m) of 19 trees among dominant species in MSH (*Fagus grandifolia*, *Acer saccharum*, *Acer pensylvanicum* and *Ostrya virginiana*). In 2018, we realized a time series survey in MSH and SBL. In each forest, we marked and collected leaf samples from the subcanopy of 40 trees (representative of local tree species diversity) in 4-6 plots distributed along a 1.2 km transect (**Supplementary Materials and Methods S1)**. In MSH, the transect followed an elevation and floristic gradient dominated by tree species *F. grandifolia* (*FAGR*), *A. saccharum* (*ACSA*), *O. virginiana (OSVI*) and *Quercus rubra* (*QURU*). In SBL, the transect followed a constant environnement dominated by *A. saccharum*, *F. grandifolia*, *A. pensylvanicum (ACPE), Abies balsamea* (*ABBA*) and *Acer rubrum* (*ACRU*). For this time series, each tree was sampled 3-4 times from June to October 2018. For each sampled plot and time point, we also sampled a negative control consisting of empty sterile bags opened and sealed on site. The leaf surface microbial community from each sample was collected with phosphate buffer and split in two equal volumes for microbial community DNA extraction and *Methylobacterium* isolation, respectively (**Supplementary Materials and Methods S1)**.

**Figure 1.**
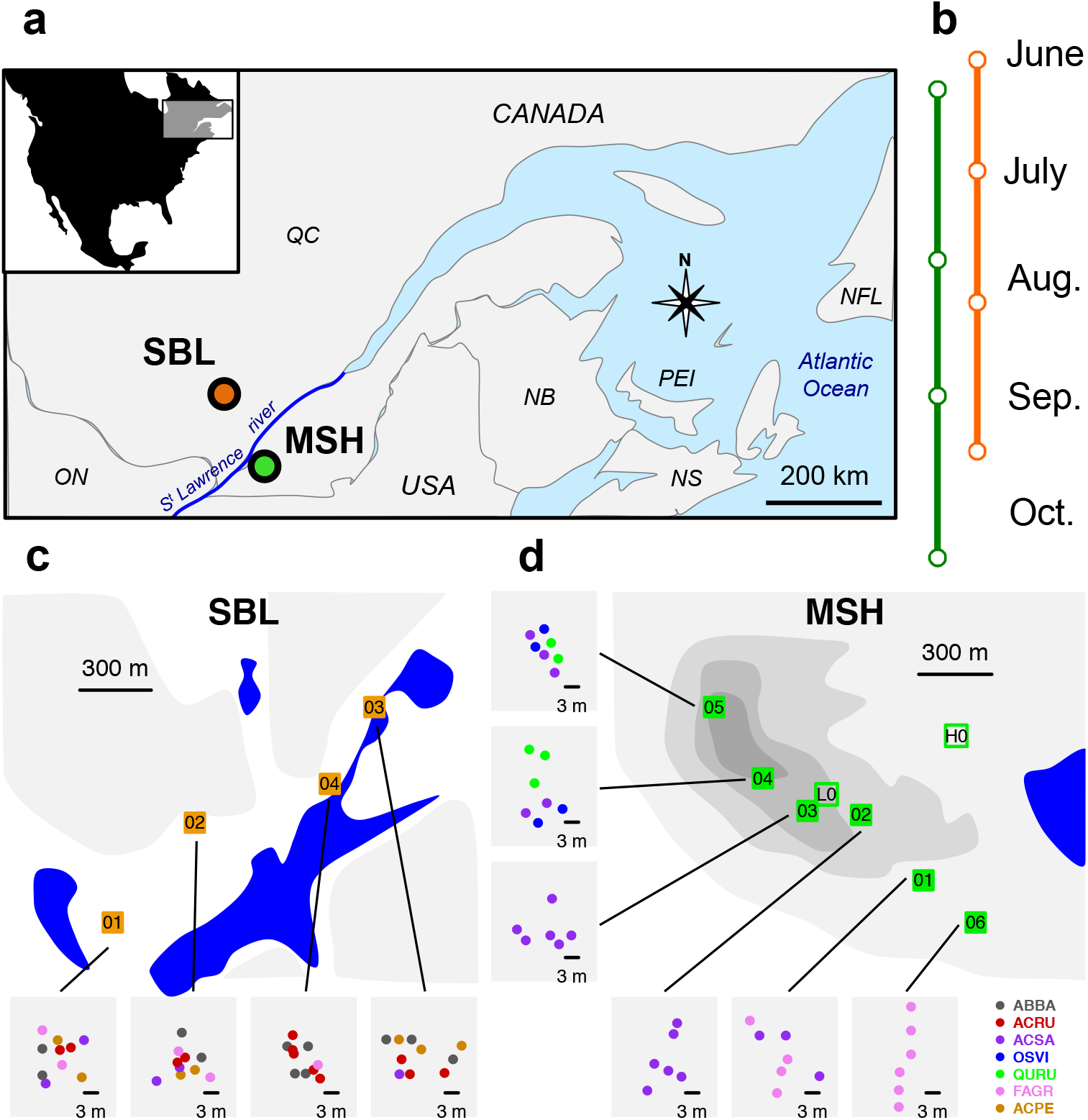
Sampling design. a) Locations of the two sampled forests MSH (green) and SBL (orange) in the province of Québec (Canada). b) Time line survey in each forest in 2018 (2-4 time points available per tree). c-d). Detailed map of each forest and each plot within forests (squares; 6 to 10 trees were sampled per plot; see **Dataset S1b**). For each plot, trees are indicated by points colored according to their taxonomy (color code on bottom left): ABBA (*Abies balsamea*), ACRU (*Acer rubrum*). ACSA (*Acer saccharum*), OSVI (*Ostrya virginiana*), QURU (*Quercus rubra*), FAGR (*Fagus grandifolia*), ASPE (*Acer Pennsylvanicum*). Shades of grey indicate elevation (50 m elevation scale)

#### Methylobacterium *isolation and development of a fine-scale single-copy molecular marker specific to Methylobacterium*

For both pilot and time series surveys, we performed *Methylobacterium* isolation on MMS synthetic solid media with 0.1% methanol supplemented with yeast extract and vitamins (**Supplementary Materials and Methods S1)**. For each leaf sample, isolation was replicated at 20 °C and 30 °C to minimize biases toward mesophylic strains. Isolates from the 2017 pilot survey (*n*=80; **Dataset S1c**) were identified by PCR amplification and partial sequencing of the *16S rRNA* ribosomal gene and assigned to *Methylobacterium* clades (**Supplementary Materials and Methods S1; Dataset S1d,e)**. As an alternative to the *16S rRNA* gene, we developed a highly polymorphic marker targeting the *Methylobacteriaceae* family. We tested two candidate genes, *rpoB* (47, 48, 50, 51) and *sucA* (51–53), which, contrary to *16S rRNA*, were single-copy in *Methylobacterium* genomes and were polymorphic enough to distinguish among *Methylobacterium* groups and clades (**Supplementary Materials and Methods S1; Figure S1)**. In 20 representative *Methylobacterium* isolates from the 2017 pilot survey (**Dataset S1c,d,e**), we successfully amplified a *rpoB* hypervariable region (targeted by primers Met02-352-F and Met02-1121-R), that we choose as a specific marker for *Methylobacteriaceae* (**Supplementary Materials and Methods S1**; **Table S1**). Isolates from the 2018 timeline survey (*n*=167; **Dataset S1e,f,**) were assigned to *Methylobacterium* clades using a consensus phylogenetic tree inferred with MrBayes v. 3.2.7a (49) from nucleotide sequences of the *rpoB* marker obtained for these isolates, aligned together with *rpoB* complete nucleotide sequences available from 188 *Methylobacteriaceae* genomes (**Dataset S1a**) and partial nucleotide sequences obtained from 20 representative isolates from the pilot survey (**Supplementary Materials and Methods S1)**.

#### *Culture-based assessment of* Methylobacterium *diversity in the tree phyllosphere*

We tested for associations between *Methylobacterium* culture-based diversity at different phylogenetic depths, with isolate characteristics as proxy for adaptive response to environmental variables through their evolution, using the *rpoB* phylogenetic tree built from timeline survey isolates as a guide. We assigned *Methylobacterium* isolates according to their phylogenetic placement. After excluding nodes supported by less than 30% of bootstraps, the tree was converted into an ultrametric tree scaled proportionally to pairwise nucleotide similarity (*PS*; **Supplementary Materials and Methods S1**). First, for each *PS* value in the tree in the range 0.926-1.000 (corresponding to *PS* range within clades), we classified isolates into discrete taxa and performed a PERMANOVA (10,000 permutations) on *Methylobacterium* community dissimilarity using the *Bray-Curtis* index (*BC*) based on taxa absolute abundance (Hellinger transformation) using the R package *vegan* (54). We tested for the relative contribution of four factors and their interactions on taxon frequency: sampling forest (*F*); temperature of isolation (*T*); sampling time (*D*) and host tree species (*H*). Second, we asked specifically which nodes within the tree were associated with *F* and *T*. For each node with at least 30% of support, and each factor, we tested for the association between embedded taxa and *F* (SBL and MSH) or *T* (20 and 30 °C) by permutation of factors between embedded nodes (100,000 permutations per node; **Supplementary Materials and Methods S1**).

#### *Culture-free assessment of* Methylobacterium *diversity in the tree phyllosphere* (*barcoding*)

We evaluated the bacterial phyllosphere diversity through barcoding and sequencing of phyllosphere samples from the 2018 timeline survey. First, we evaluated the bacterial diversity targeting the *16S rRNA* gene (55) in 46 phyllosphere samples from 13 trees from both forests sampled 3-4 times throughout the 2018 growth season. We included one negative control, and one positive control consisting of mixed DNAs of *Methylobacterium* isolates typical of the phyllosphere (METH community; **Supplementary Materials and Methods S1**). Second, we evaluated the *Methylobacteriaceae* phyllosphere diversity targeting the *rpoB* marker (see above) in 184 phyllosphere samples from 53 trees representative of diversity found in MSH (n=26) and SBL (n=27), sampled 3-4 times throughout the 2018 growth season. We included four negative controls and four positive controls (METH community). Library preparation and sequencing were performed as described in **Supplementary Materials and Methods S1.** For each phyllosphere sample and controls, we estimated bacterial diversity based on Amplicon Sequence Variants (ASVs) using package *dada2* in R (56). We assessed ASV taxonomy using SILVA v.138 database for *16S rRNA* gene (57) and a *rpoB* nucleotide sequence database available for *Bacteria* (48), curated by a ML phylogenetic tree (200 permutations**; Supplementary Materials and Methods S1**). Taxonomy for *Methylobacterium* ASVs (at the clade level) was refined using blast against NCBI databases for *16S rRNA* gene (58) and using phylogenetic placement for *rpoB* (**Supplementary Materials and Methods S1**). To validate the *rpoB* barcoding accuracy in estimating *Methylobacterium* diversity, we compared *Methylobacterium* clade relative abundances estimated from *16S rRNA* and *rpoB* barcoding in a heatmap (**Supplementary Materials and Methods S1**). We also compared *Methylobacterium* diversity estimations from *rpoB* barcoding and culture-dependant approaches by matching *rpoB* partial nucleotide sequences obtained from isolates with those obtained from ASVs (**Supplementary Materials and Methods S1**). We evaluated relative contributions of sampling forest (*F*), plot within forest (*P*), host tree species (*H*), time of sampling (*T*) and their interactions on bacteria (*16S rRNA* barcoding) and *Methylobacteriaceae (rpoB* barcoding) community dissimilarity among phyllosphere samples (*BC* index, Hellinger transformation on ASV relative abundance), using PERMANOVA (10,000 permutations; **Supplementary Materials and Methods S1**) and principal component analysis (PCA). For *rpoB* barcoding, specifically, we reported *Methylobacterium* ASV significantly associated with the aforementioned factors (*F*, *P*, *H*, *T*; ANOVA) into the PCA (**Supplementary Materials and Methods S1)**.

#### *Spatial and temporal dynamics of* Methylobacterium *communities*

We evaluated the spatial and temporal dynamics of *Methylobacterium* communities in the timeline survey (*rpoB* barcoding) using autocorrelation analyses. In order to remove potential differences in community composition between forests, we analyzed samples from MSH and SBL separately. For each pairwise comparison between two samples from the same forest, we evaluated the effects of spatial distance (*pDist*) separating trees sampled at the same date (*spatial autocorrelation analyses*) and time (*pTime*) separating dates at which trees were sampled (*temporal autocorrelation analyses*) on *BC* dissimilarity among samples (see above). We evaluated the effects of *pDist* and *pTime* on *BC* under linear models by ANOVA (**Supplementary Materials and Methods S1**).

#### *Ecophylogenetic structure of* Methylobacterium *communities*

We quantified the ecophylogenetic structure of *Methylobacterium* communities by comparing the phylogenetic dissimilarity of co-occurring *rpoB* ASVs with the dissimilarity expected under a null model of stochastic community assembly from the pool of all ASVs, in order to quantify the evidence for different community assembly processes (59) as a function of forest, host tree species, and time of sampling. For each community of *Methylobacterium* ASVs, we calculated a measure of phylogenetic dissimilarity among co-occurring ASVs (mean nearest taxon distance (MNTD)) and compared observed MNTD to that expected under a null model of stochastic community assembly from the pool of all ASVs. We calculated the standardized effect size (SES) of MNTD (60), which expresses the difference between the observed MNTD value versus the mean and standard deviation of MNTD values obtained across 999 random draws of ASVs from the pool of observed ASVs across all samples while maintaining observed sample ASV richness (61). We evaluated the effects of forest, host tree species, and time of sampling on SES(MNTD) by ANOVA.

#### *Monitoring of* Methylobacterium *growth performance*

We evaluated the growth abilities of 79 *Methylobacterium* isolates from the timeline survey for four temperature treatments mimicking temperature variations during the growing season. Each treatment consisted of an initial pre-conditioning step (*P*) during which each isolate was incubated on solid MMS media with methanol as sole carbon source for 20 days at either 20 °C (*P20*) or 30 °C (*P30*), and a second monitoring step (*M*) during which pre-conditioned isolates were incubated on the same media and their growth monitored for 24 days at 20 °C (*P20M20* and *P20M30*) or 30 °C (*P30M20* and *P30M30*; **Figure S2**) Treatments *P20M20* and *P30M30* mimicked stable thermal environments, and treatments *P20M30* and *P30M20* mimicked variable thermal environments. For each isolate and each combination of treatments (*PXXMXX*), we realized 5 replicates, randomly spotted on 48 petri dishes according to a 6×6 grid. During the monitoring step, we took photographs of each petri dish at days 7, 13 and 24 after inoculation (**Figure S2**). Photos were converted to pixel intensities with ImageJ 1.52e and processed in R for background correction, measurement of spot intensities and correction for position-dependant competition effects (**Figure S3**; **Supplementary Materials and Methods S1).** For each isolate and temperature treatment, logistic growth curves were inferred from bacteria spot intensity variation observed over three time points during the monitoring step. From growth curves, we estimated maximum growth intensity, or yield (*Y*) and growth rate (*r*) as the inverse of lag+log time necessary to reach *Y* (**Figure S4** (62, 63); **Supplementary Materials and Methods S1**). We evaluated the effects of following factors on *Methylobacterium* growth abilities (*Y* and *r*) under different temperature treatments: isolate assignement to clades (*C*), forest of origin (*F*), host tree species (*H*), time of sampling (*D*), temperature of isolation (*T_I_*; at which each isolate was isolated), temperature of incubation during pre-conditioning (*T_P_*) and monitoring (*T_M_*) steps, and all possible interactions between those factors (ANOVA; **Supplementary Materials and Methods S1**).

## Results

### Phylogenetics of plant-associated Methylobacterium diversity

A phylogeny of 153 *Methylobacterium* isolates built from available genomic databases showed that plants (65% of strains) and especially the phyllosphere compartment (41% of strains) were the most prevalent source of *Methylobacterium* sampled to date (**Figure 2; Dataset S1a**). Phyllosphere-associated diversity was not randomly distributed in the *Methylobacterium* phylogenetic tree. Isolates from the phyllosphere represented the largest part of diversity within group A (56% of isolates) but not in groups B and C (17 and 12% of isolates, respectively). Group A was paraphyletic and most of its diversity consisted of undescribed taxa falling outside previously well-described linages. Accordingly, we subdivided *Methylobacterium* group A into 9 monophyletic clades (A1-A9).

**Figure 2.**
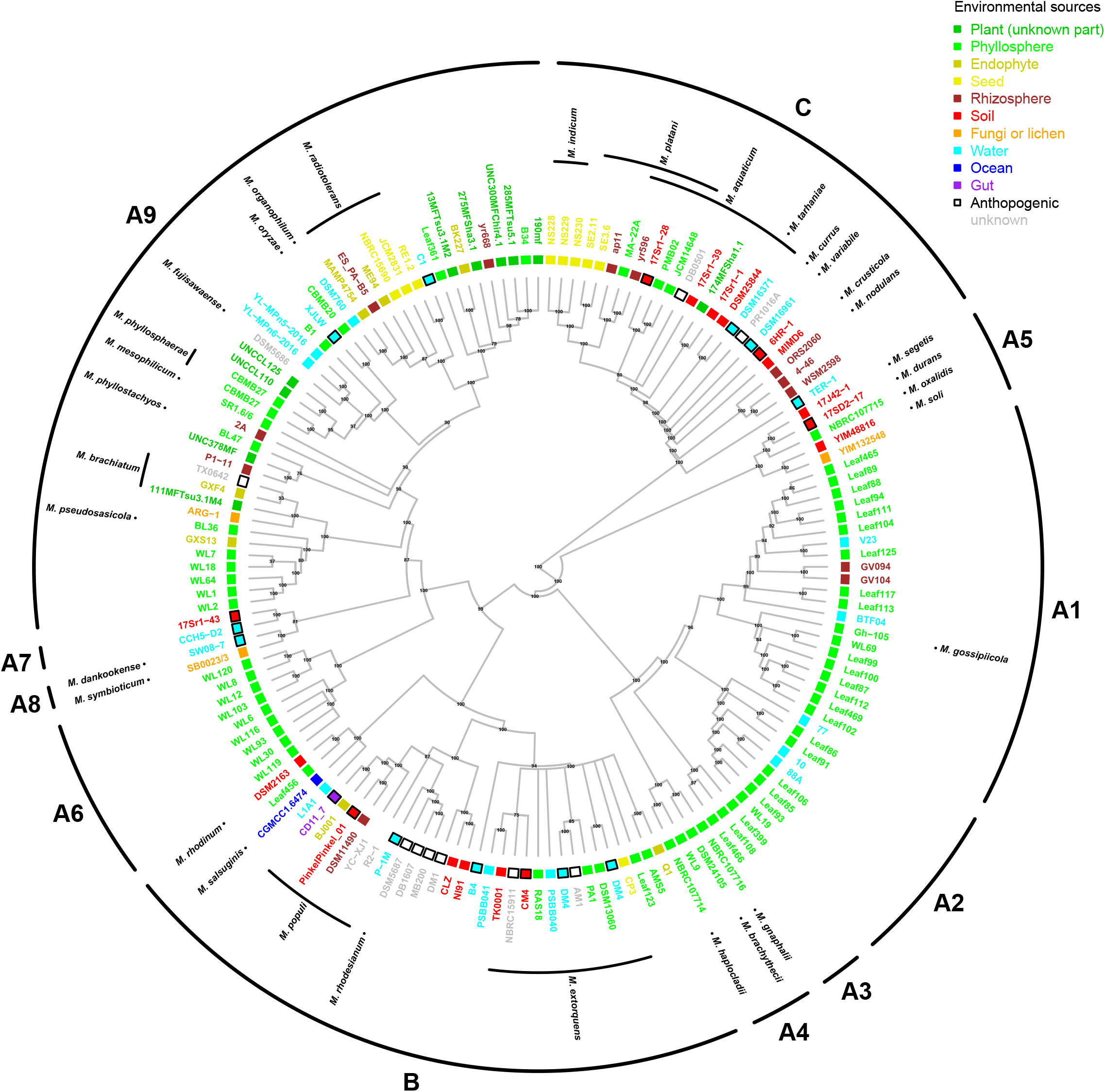
*Methylobacterium* phylogeny and ecology. Most of *Methylobacterium* diversity is found in association with plants, especially in the phyllosphere. Phylogenetic consensus tree (nodal posterior probabilities indicated next to the branches) from *rpoB* complete nucleotide sequences available for 153 *Methylobacterium* genomes and rooted on 32 *Methylobacteriaceae* outgroups (*Microvirga, Enterovirga*; no shown; see **Dataset S1a**). For each genome, species name, the anthropogenic origin (black squares) and/or environmental origin (color code on top right) are indicated. Groups A, B, C adapted from Green *et* Ardley (31).

#### 16S rRNA *community analyses of the tree phyllosphere*

We focused on *Methylobacterium* phyllosphere diversity variation observable at the scale of seasonal variation (within year 2018) on individual trees within two temperate forests of northeastern North America (**Figure 1a,b; Dataset S1b,g**): Mont Saint Hilaire (MSH**; Figure 1c**) and Station biologique des Laurentides (SBL; **Figure 1d**). The distribution of the phyllosphere bacterial community assessed in 46 leal samples by bacterial *16S rRNA* gene amplicon sequence variants (ASVs) was mostly explained by differences among forests (31.6% of variation explained; p<0.001; PERMANOVA), host tree species (15.6% of variation; p<0.001) and time of sampling (12.0%; p<0.05; **Table 1**). Although representing only 1.3% (0.0-3.2% per sample) of total *16S rRNA* sequence diversity, *Methylobacterium* was present in almost all analyzed samples (45 out of 46; **Dataset S1h**). We assigned the 15 *Methylobacterium* ASVs identified by *16S rRNA* sequencing to clades from *Methylobacterium* group A: A9 (related to *M. phyllosphaerae/M. mesophilicum/M. phyllostachyos/ M. pseudosasicola/M. organophilum*; 0.87% of total diversity, nine ASVs), A6 (related to *M. cerastii*, 0.29%; one ASV) and A1 (related to *M. gossipicola*; 0.13%, 3 ASVs; **Table S2**; **Dataset S1i**). With two rare ASVs (<0.01% of relative abundance) related to *M. komagatae*, belonging to group A (31) but unrelated to any aforementioned clade, we defined a new clade (A10). No ASVs from MSH or SBL were assigned to group B or group C.

**Table 1.** PERMANOVA analysis of variance in *Bacteria* and *Methylobacterium* community diversity. Part of variance in dissimilarity (*R^2^*; Bray-Curtis index) among samples associated with four factors and their possible interactions (*F*: forest of origin; *D*: date of sampling; *H*: host tree species; *P*: plot within forest) and their significance are shown (10,000 permutations on ASV relative abundance, Hellinger transformation; “***”: p<0.00l; “**”: p<0.01; “*”: p<0.05.). For *16S rRNA*, *P* was omitted to conserve degrees of freedom.

|  | *Bacteria* (16S rRNA) |  | *Methylobacterium* (*rpoB*) |  |
| --- | --- | --- | --- | --- |
| Samples | 46 |  | 179 |  |
| Factor | $R^2$ | Pr(>F) | $R^2$ | Pr(>F) |
| Forest of origin (F) | 0.316*** | <0.000 | 0.324*** | <0.001 |
| Host tree specie (H) | 0.156*** | <0.001 | 0.071*** | <0.001 |
| Time of sampling (D) | 0.120* | 0.016 | 0.048*** | <0.001 |
| Plot within forests (P) | - | - | 0.080*** | <0.001 |
| F:H | 0.020 | 0.080 | 0.004 | 0.110 |
| H:D | 0.239 | 0.217 | 0.074** | 0.028 |
| H:P | - | - | 0.043** | 0.007 |
| D:P | - | - | 0.058 | 0.455 |
| H:D:P | - | - | 0.085 | 0.052 |
| Residuals | 0.150 | - | 0.213 | - |

### Culture-based assessment of Methylobacterium diversity in the tree phyllosphere

We evaluated the culturable part of *Methylobacterium* diversity from a subsample of 36 trees (18 per forest). Using *rpoB* gene partial nucleotide sequences as a marker, we identified 167 pink isolates that we assigned to *Methylobacterium* based upon their phylogenetic placement (**Dataset S1e,f; Figure 3)**. As observed for *16S rRNA* ASVs, most isolates were assigned to clades from group A typical of the phyllosphere: A9 (59.9% of isolates), A6 (24.6%), A1 (5.4%), A10 (3.6%) and A2 (related to *M. bullatum* and *M. marchantiae*; **Dataset S1d** 1.8%). Few isolates were assigned to group B (4.2% of isolates, related to *M. extorquens*) and none to group C (**Table S2**). The higher polymorphism in the *rpoB* marker revealed a considerable diversity within clades, as we identified 71 unique *rpoB* sequences, in contrast to the smaller number obtained with *16S rRNA* barcoding (15 ASVs). We determined that *Methylobacterium* diversity assessed at varying depths in the *rpoB* phylogeny was systematically explained by forest of origin (4.5±1.0% of variance explained; PERMANOVA; *p*<0.001; **Figure 3a; Dataset S1j**) and temperature of isolation (5.9±2.1% of variance explained; *p*<0.001). Temperature of isolation was the most important factor distinguishing deep phylogenetic divergences (pairwise nucleotide similarity range: 0.948-0.993), while forest of origin was slightly more important in structuring more recently diverged nodes (pairwise nucleotide similarity >0.993). Time of sampling had a slight but significant effect on diversity (2.1±0.2% of variance explained; *p*<0.05) and it was only observed for higher pairwise nucleotide similarity values (range 0.994-1.000). We did not observe any significant effects of host tree species on *Methylobacterium* isolate diversity, at any level of the phylogeny. In the phylogeny, we identified two nodes strongly associated with temperature of isolation, corresponding to clades A6 (20 °C; *p*<0.001; permutation test) and A9+A10 (30 °C; *p*<0.001; **Figure 3b**). Other clades were evenly isolated at 20 and 30 °C and we observed no significant association between temperature of isolation and nodes embedded within clades. Nodes associated with forest of origin also roughly corresponded to certain major clades, with clades A1+A2 almost exclusively sampled in MSH (*p*<0.01). Overall, clade A9 was isolated significantly more often at SBL (*p*<0.001) but at least three of its subclades were significantly associated with either MSH or SBL (*p*<0.05).

**Figure 3.**
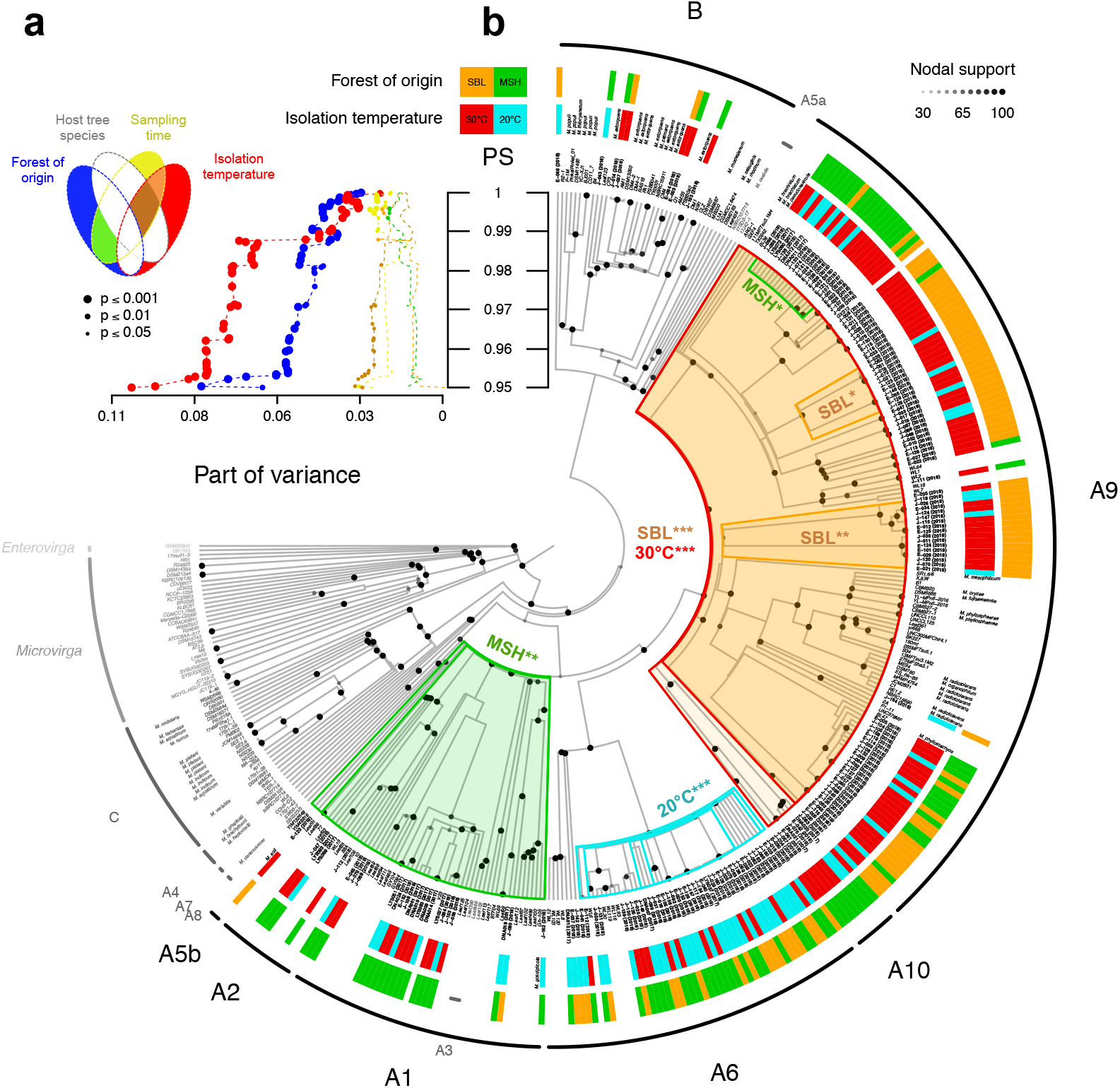
Tests for phylogenetic association of traits with culture-based estimation of *Methylobacterium* diversity. a) Part of variance (PERMANOVA; x-axis) in *Methylobacterium* isolated diversity explained by forest of origin, host tree species, sampling date, temperature of isolation and their interactions (see Venn diagram on top left for color code) in function of pairwise nucleotide similarity (PS; y-axis; see **Dataset S1j**) in a phylogenetic tree (partial *rpoB* nucleotide sequences of 187 isolates and 188 *Methylobacteriaceae* reference sequences). b) Permutation test for node association with forest of origin and temperature of isolation (color code on top) mapped on the *rpoB* phylogeny (scaled on PS values). Frames in the tree indicate nodes significantly associated with at least one factor (ANOVA; Bonferroni correction; *p*<0.001: “***”: *p*<0.01 *p*<0.05:”*”). For each isolate (names in bold), colored boxes at the tip of the tree indicate forest of origin and temperature of isolation.

### Comparison of Methylobacterium diversity assessed by rpoB barcoding and isolation

We performed culture-independent *rpoB* amplicon sequencing from 179 leaf samples from 53 trees in both forests, allowing a monthly monitoring for most trees (**Dataset S1b,g**). We identified 283 *Methylobacteriaceae rpoB* ASVs in these samples (**Dataset S1k,l**), representing 24.6% of all sequences. *Non-Methylobacteriaceae* ASVs were mostly assigned to other Rhizobiales families (850 ASVs, 70.33% of sequence abundance) and to Caulobacterales (209 ASVs, 4.42% of sequence abundance) typical of the phyllosphere (**Supplementary Materials and Methods S1**), indicating that the *rpoB* marker can potentially be used at a broader taxonomic scale (**Figure S5a**). Within *Methylobacteriaceae*, ASVs were mostly classified as *Methylobacterium* (200 ASVs, 23.05% of sequence relative abundance), and *Enterovirga* (78 ASVs, 1.56%; **Dataset S1k**). We assigned most of *Methylobacterium* ASVs to previously cultured clades A9 (45.2% of *Methylobacterium* sequence abundance), A6 (24.3%), A1 (6.1%) and A10 (1.0%; **Dataset S1k; Table S2; Figure S5b**). Estimates of *Methylobacterium* diversity based on *rpoB* sequences from culture-independent sequencing were generally concordant with estimates based on *16S rRNA* barcoding (**Figure S5c; Table S2**) and estimates from cultured isolates (**Figure S5d; Table S2**). The major exception was group B, representing 19.1% of *Methylobacterium* sequence abundance (*rpoB* barcoding) but not detected by *16S rRNA* barcoding, and representing 4.2% of isolates (**Table S2**). Clade A4 (related to *M. gnaphalii* and *M. brachytecii*) represented 1.7% of *Methylobacterium* sequence abundance (*rpoB* barcoding) but was not detected by *16S rRNA* barcoding, nor was it isolated. Other clades could be detected by *rpoB* barcoding with low sequence abundance (<0.3%) but not by *16S rRNA* barcoding, and were unevenly isolated (<1.8% of isolates).

### Fine-scale temporal and spatial distribution of Methylobacterium diversity assessed by rpoB barcoding

The community composition of the 200 *Methylobacterium* ASVs was mostly explained by spatial variation at both large (distance between forests: 100 km) and local scales (distance between plots within forest: 150-1,200 m), as well as sampling date during the growing season (1-5 months; proportion of variation explained: 32.4%, 8.0% and 4.8%, respectively; p<0.001; PERMANOVA; **Table 1**). We observed slight but significant effects of host tree species, and of the interaction between host tree species and plots within forests, on *Methylobacterium* community composition (explaining 7.1% and 4.3% of variation in community composition; p<0.001 and p<0.01, respectively; PERMANOVA; **Table 1**). A large proportion of *Methylobacterium* ASVs (83 out of 200) were significantly associated with one or either forest (ANOVA; **Figure 4a; Dataset S1m**), regardless their clade membership. The only exception was clade A1, which was almost exclusively observed (and isolated; see **Figure 3b**) in the MSH forest. We found 25 ASVs whose relative abundance significantly increased throughout the growing season (ANOVA; p<0.05), mostly belonging to clades A1 (n=11). Four ASVs increased significantly in frequency over time in both forests, and mostly belonged to group B (n=3), (**Dataset S1m**). We found no clear association between ASV or clade with host tree species, nor plots within forests (data not shown). *Methylobacterium* diversity was heterogeneously distributed at local spatial scale, as we observed a significant increase of community dissimilarity (Bray-Curtis index; *BC*) with geographical distance separating two samples within MSH (spatial autocorrelation analysis; ANOVA; *p*<0.001) but not SBL (*p*>0.05, **Table 2, Figure 4b**). We also observed a significant increase of community dissimilarity over time separating two sampling dates in both forests (temporal autocorrelation analysis; *p*<0.001; **Table 2**), indicating that community composition changed during the growing season. This effect was more marked in MSH than in SBL (**Figure 4c**). The overall community *BC* dissimilarity consistently decreased from June to October in both MSH (from 0.624 to 0.297) and SBL (from 0.687 to 0.522; **Table 2, Figure 4d**), indicating that the observed change of diversity over time resulted from a progressive homogeinization of *Methylobacterium* community between the beginning and the end of the growing season at the scale of a forest, although without affecting its heterogeneous spatial distributions in MSH (**Table 2, Figure 4e**). *Methylobacterium* communities were strongly phylogenetically clustered (**Figure 4f**), with all communities containing ASVs that were much more closely related than expected by chance (mean SES(MNTD) (± standard deviation) = −4.8 ± 0.9, all SES(MNTD) *p*-values <0.05 compared with null model of random community assembly). While all communities were strongly phylogenetically clustered, SES(MNTD) differed among host tree species (ANOVA; F=6.4, *p*<0.001) and forests (ANOVA; F=10.9, P<0.001), and decreased during the growing season (ANOVA; F=95.2, *p*<0.001).

**Figure 4.**
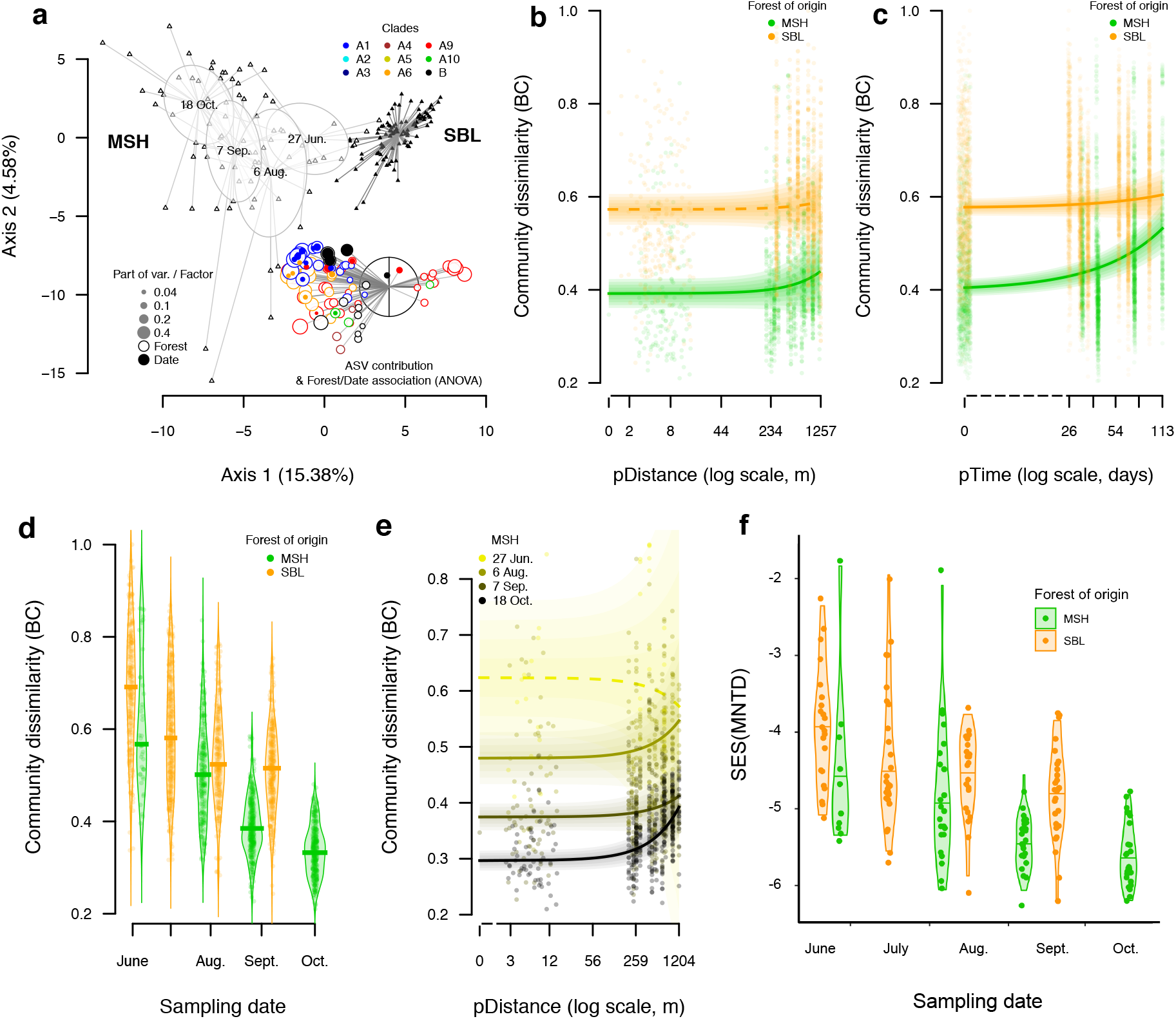
Short-scale spatial and temporal dynamics of *Methylobacterium* communities assessed by *rpoB* barcoding. a) A principal component analysis (PCA) on *Methylobacterium* ASVs relative abundance shows that 179 phyllosphere samples cluster according to forest of origin (MSH: open triangles, SBL: full triangles) and date of sampling (detail showed only for MSH). The significant association of 83 and 25 ASVs with forest of origin and/or sampling date, respectively is shown (points colored according to clade assignation; legend on top right). b) Spatial and c) temporal autocorrelation analyses conducted in each forest separately. Points represent Bray-Curtis (*BC*) dissimilarity in function of pairwise geographic (*pDist*; b) or pairwise time (*pTime*; c) distance separating two communities. For each forest and variable, the predicted linear regression is indicated (full line: *p*<0.001; dotted line: *p*>0.05; ANOVA). d) *BC* in function of sampling time for each forest. e) Detail of spatial autocorrelation analyzes in MSH, conducted for each sampling time point separately. f) Standardized effect size of mean nearest taxon phylogenetic distance (SES(MNTD)) between forests and across sampling dates. Negative values of SES(MNTD) indicate communities contain ASVs that are phylogenetically clustered compared to a null model of stochastic community assembly.

**Table 2.** Summary of statistics from autocorrelation analyzes on 179 phyllosphere *Methylobacterium* samples assessed by *rpoB* barcoding (200 ASVs). *Spatial autocorrelation general models*: pairwise dissimilarity between two communities (Bray-Curtis index; *BC*) as a function of pairwise spatial distance separating two sampled trees (*pDist*) and date of sampling (*Date*) and their interaction (*pDist:Date*). *Spatial autocorrelation models per date: BC* as a function of pairwise spatial distance (*pDist*). *Temporal autocorrelation: general models: BC* as a function of pairwise spatial time separating two sampled trees (*pTime*). For each model, the average and standard deviation of the intercept (mean *BC* value) are indicated. For each factor (*pDist*, *Date*, *pDist:Date* and *pTime*), the average and standard deviation of estimates (slope) are indicated. Significance of estimates was assessed by ANOVA (“***”: p<0.00l; “**”: p<0.01; “*”: p<0.05).

| Categories (n) | Intercept (sd) |  | Estimates*10<sup>-3</sup> (sd) |  |
| --- | --- | --- | --- | --- |
| Spatial autocorrelation general models: lm(BC~pDist*D) |  |  |  |  |
| Site (within dates) | BC | pDist | D | pDist:Date |
| MSH | 0.5965 (0.0107) | -0.0041 (0.0192)*** | -2.7648 (0.1313)*** | 0.0007 (0.0002)** |
| SBL | 0.6493 (0.0097) | 0.0157 (0.0145) | -1.5575 (0.1646)*** | 0.0000 (0.0002) |
| Spatial autocorrelation models per date: lm(BC~pDist) |  |  |  |  |
| Site | Date | BC | pDist |  |
| MSH | 27 Jun. | 0.6237 (0.0340) | -0.0425 (0.0725) |  |
|  | 6 Aug. | 0.4919 (0.0112) | 0.0503 (0.0192)** |  |
|  | 7 Sept. | 0.3746 (0.0059) | 0.0313 (0.0099)** |  |
|  | 18 Oct. | 0.2966 (0.0045) | 0.0795 (0.0073)*** |  |
| SBL | 20 Jun. | 0.6868 (0.0146) | 0.0082 (0.0216) |  |
|  | 16 Jul. | 0.5819 (0.0113) | 0.0215 (0.0174) |  |
|  | 16 Aug. | 0.5415 (0.0105) | 0.0114 (0.0150) |  |
|  | 20 Sept. | 0.5222 (0.0089) | 0.0145 (0.0130) |  |
| Temporal autocorrelation general models (BC~pTime) |  |  |  |  |
| Site | BC | pTime |  |  |
| MSH | 0.4086 (0.0032) | 1.0786 (0.0607)*** |  |  |
| SBL | 0.5789 (0.0030) | 0.3012 (0.0617)*** |  |  |

### Effect of short scale temperature variation in combination with other environmental and genetic factors on Methylobacterium growth performances

We measured growth of 79 *Methylobacterium* isolates (sampled in 2018 in both forests; MSH: *n*=32, SBL: *n*=47) in conditions mimicking temperature variations during the growing season (**Figures S2-S4**; **Dataset S1n**). Clade membership explained a large part of variation in growth rate (*r*) and yield (*Y*; 7.6 and 30.6% of variation explained, respectively; ANOVA; *p*<0.001; **Figures 5a,b, Table 3; Dataset S1o**). Group B isolates (*Y* = 12.2 ± 5.0) have higher yield than group A (*Y* = 5.4 ± 3.5). Isolates from clades A1, A2 and B had the highest growth rate (*r* range: 0.101±0.032 – 0.121±0.031). Other clades (A6, A9 and A10) had on average slower growth (*r* range: 0.082±0.021 – 0.088±0.024). Time of sampling, host tree species and forest also explained significant variation in growth rate (5.4%, *p*<0.001; 2.2%, *p*<0.01 and 1.5%, *p*<0.05, respectively; ANOVA) and limited or no significant variation in yield (1.3%; *p*<0.001; 1.3%; *p*<0.01; 0.2%; *p*>0.05, respectively; **Table 3**). Among the aforementioned factors only the interaction between time of sampling and clade membership explained significant variation in growth rate (2.9%; *p*<0.001), while all possible pairwise interactions between these factors explained significant variation in yield (range 1.4 – 5.9%; *p*<0.01; **Table 3**). In both SBL and MSH, growth rate increased consistently from June (*r* = 0.075±0.018 and 0.085±0.033, respectively) to September/October (*r* = 0.097±0.031 and 0.103±0.027, respectively; **Figure 5c**). The temperature of isolation (at which each isolate was originally isolated) had very limited effect on growth rate (1.0%; *p*<0.01) and yield (0.6%; *p*<0.05). These effects were independent of temperatures during pre-conditioning and monitoring steps (no significant interaction in the ANOVA). Temperature of incubation had significant effects on growth performance. Temperature during the monitoring step explained respectively 2.0% and 15.8% of variation in yield and growth rate (*p*<0.01 and *p*<0.001, respectively; ANOVA; **Table 3**), regardless of clade membership, time of sampling, and other environmental factors (no significant interaction in the ANOVA). Isolates incubated at 20 °C had on average higher yield (*Y*=6.9±5.4) but slower growth (*r*=0.077±0.022) than isolates incubated at 30 °C (*Y*=4.9±3.6; *r*=0.100±0.030; **Figure 5d**). Temperature during the pre-conditioning step had no effect on growth rate (*p*>0.05; ANOVA), and limited effect on yield (1.4%; *p*<0.05; ANOVA; **Table 3**).

**Figure 5.**
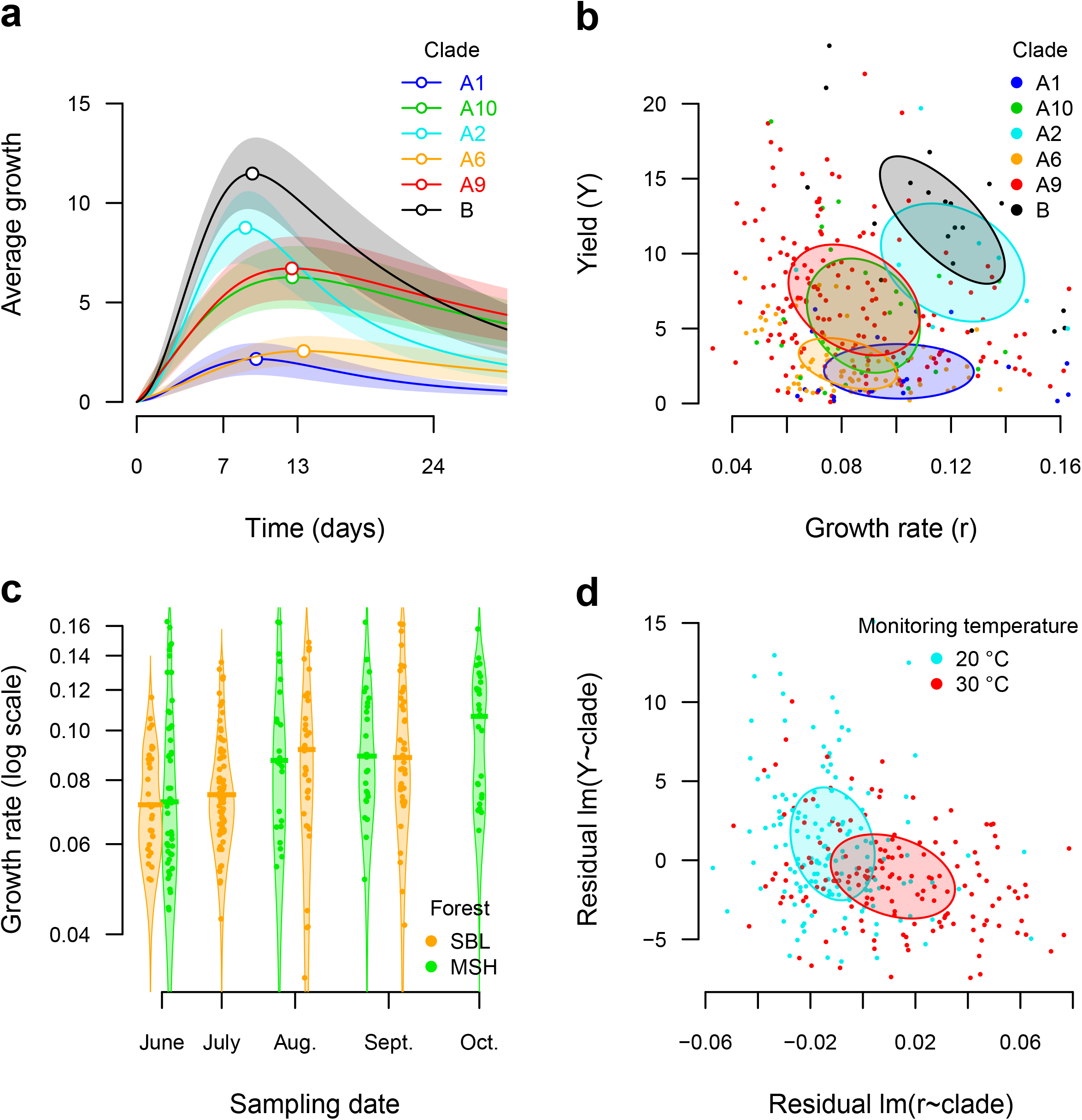
Analysis of 79 *Methylobacterium* isolate growth performances under 4 different temperature treatments. a) Average growth curves (growth intensity in function of time) for each clade (line: mean value; frame: 1/3 of standard deviation; point: average maximal growth). b) Growth rate (*r*) in function of yiel (*Y*). Each point represents the average *r*/*Y* values for an isolate and a temperature treatment (79 isolates x 4 treatments), colored according to clade membership. Ellipsoides are centered on average values per clade and represent 30% of confidence interval (standard deviation). c) *r* (log scale) in function of time at which samples strains were isolated from were collected, colored according to the forest of origin. Points: real data; bars: average *r* value per forest (n=2) and time (n=4) category. d) *r* in function of Y, corrected for clade assignement (residuals of the *r*~Clade and *Y*~Clade linear regressions). Each point represents the average *r*/*Y* residual values for an isolate and a temperature, colored according to the monitoring temperature (legend on top right).

**Table 3.** Variance in yield (*Y*) and growth rate (*r*) measured in 79 *Methylobacterium* isolates grown under 4 temperature treatments. *Y* and *r* values were transformed in log to meet normal distribution. For each factor following factors: clade (*C*), forest of origin (*F*), host tree species (*H*), time of sampling (*D*), temperature of incubation during pre-conditioning (*T_B_*) and monitoring (*T_M_*) steps, temperature of isolation (*T_I_*) and their interactions, significance of *Y* and *r* responses are shown (“***”: p<0.00l; “**”: p<0.01; “*”: p<0.05; see **Dataset S1o** for details).

|  | Rate | Yield |
| --- | --- | --- |
| Forest ($F$) | 0.015** | 0.002 |
| Host tree species ($H$) | 0.022*** | 0.013*** |
| Date of sampling ($D$) | 0.054*** | 0.013*** |
| Pre-conditioning temperature ($T_P$) | 0.001 | 0.014*** |
| Monitoring temperature ($T_M$) | 0.158*** | 0.020*** |
| Temperature of isolation ($T_I$) | 0.010** | 0.006* |
| Clade ($C$) | 0.076*** | 0.306*** |
| $F:D$ | 0.005 | 0.035*** |
| $H:D$ | 0.003 | 0.019*** |
| $H:C$ | 0.005 | 0.059*** |
| $D:C$ | 0.029** | 0.028*** |
| $F:C$ | 0.011 | 0.014** |
| $F:T_I$ | 0.003 | 0.021*** |
| $H:T_I$ | 0.000 | 0.021*** |
| $C:T_I$ | 0.024*** | 0.007 |
| $F:H:D$ | 0.012** | 0.023*** |
| $F:H:C$ | 0.010** | 0.002 |
| $F:D:C$ | 0.001 | 0.008** |
| $H:D:C$ | 0.000 | 0.013*** |
| $H:D:T_I$ | 0.016*** | 0.019*** |
| other interactions (sum) | 0.177 | 0.079 |
| Residuals | 0.371 | 0.279 |

## Discussion

*Methylobacterium* is ubiquitous on leaves in the temperate forests of Québec and its diversity in this habitat is quite similar to what has been described in the phyllosphere throughout the world, with three main clades A9 (*M. brachiatum*, *M. pseudosasicola*), A6 (related to *M. cerastii*) and A1 (related to *M. gossipicola*) dominating diversity. Our barcoding approach based on a clade-specific *rpoB* marker revealed previously undocumented diversity within these clades, as well as within several other clades that were not detected by a classical *16S rRNA* marker: B (related to *M. extorquens*), A2 (related to *M. bullatum* and *M. marchantiae*), A4 (related to *M. gnaphalii* and *M. brachytecii*) and A10 (related to *M. komagatae*). This diversity, like that of the overall phyllosphere community, was mostly determined by differences between forests, with barcoding approaches suggesting combined effects of restricted migration, local adaptation to host tree species, and climatic conditions at large geographical scales (>100km). With higher molecular resolution, we observed that *Methylobacterium* diversity was spatially structured even at the scale of a forest (within 1.2 km), and also showed a clear pattern of temporal dynamics and succession over the course of a growing season. This result indicates that, although representing a stable proportion of the plant leaf microbiota between years (22) *Methylobacterium* diversity is highly dynamic within the course of a season. A finer analysis of *Methylobacterium* diversity suggested that clade identity partly explained *Methylobacterium* geographical distribution at large scales (between forests) but not at finer scales (plots), nor was it an indicator of adaptation to a particular host tree species, nor a determinant of temporal dynamics. These results are consistent with previous observations that geographic origin is a stronger driver of phyllosphere *Methylobacterium* diversity than host identity (22). The distribution of *Methylobacterium* diversity at small temporal and geographical scales likely resulted from more contemporaneous community assembly events selecting for phenotypic traits that evolved among deeply diverging lineages of *Methylobacterium*, as has been observed in other bacterial (16) and plant clades (64). We found further evidence for deterministic community assembly as *Methylobacterium* communities were strongly phylogenetically clustered compared to the expectation under a stochastic model of community assembly, indicating that the leaf habitat acts as an ecological filter selecting for a non-random subset of *Methylobacterium* diversity.

We explored mechanisms explaining the temporal dynamics of *Methylobacterium* diversity at the scale of a growing season. Because we observed contrasting *Methylobacterium* culturable diversity between 20 and 30 °C, we suspected that adaptation to temperature variation during the growing season could explain part of these temporal dynamics. By monitoring *Methylobacterium* isolate growth under different temperature treatments, we confirmed that temperature affected isolate growth performances but interestingly, independantly from the temperature at which isolates were obtained. The fact that most tested isolates also grew slower but more efficiently at 20 °C than at 30 °C (**Figure 5d**), regardless of their phylogenetic and environmental characteristics, is in line with a temperature-dependent trade-off between growth rate and yield described in many bacteria (reviewed in (63)). High yield strategies are typical of cooperative bacterial populations, while fast growth-strategies are typical of competitive populations (63). These observations also stress the importance of considering incubation temperature when interpreting results from previous culture-based assessments of *Methylobacterium* diversity.

We provide two lines of evidence that factors other than direct adaptation to temperature drive *Methylobacterium* responses to temperature variation, by affecting their growth strategy in different competitive conditions rather than by affecting their metabolism directly. First, clade identity was one of the main predictors of overall isolate performance, with some clades (A1, A2, B) possessing a rapid growth strategy under all temperature conditions, while others (clades A6, A9, A10) had systematically slower growth. These clade-specific growth strategies could explain why certain *Methylobacterium* isolates are less competitive and less frequently isolated at higher temperatures. Still, we cannot rule out that clade-specific growth strategy also reflect experimental conditions. Second, we observed strong associations between isolate growth performance and time of sampling, regardless of clade membership, suggesting that growth strategies also respond to seasonal variations in environmental conditions, and to the level of establishment and competition in the phyllosphere community (63). These associations are unlikely to be driven by the direct effects of temperature on metabolic rates because isolation temperature had little effect on growth strategies, in contrast to clade identity and time of sampling which had more significant effects. Together, these observations could explain why isolates from clades A1 and B with fast-growth strategies consistently increase in frequency during this period due to changes in selection for different ecological strategies, leading to the homogeneization of the community.

Taken together, our temporal survey of diversity dynamics and screening for growth performance suggest the following timeline of the dynamics of the *Methylobacterium* phyllosphere community. At the very beginning of the growing season, a pool of bacteria with mixed ecological strategies and genotypes colonizes newly emerging leaves. Due to the stochasticity of this colonization, we initially observe strong dissimilarity among phyllosphere communities, regardless of their spatial position. During the summer, conditions allow the progressive establishment of a diverse *Methylobacterium* community with a high yield strategies (63), dominated by increasingly closely phylogenetically related strains. At the end of the growing season, with migration, environmental conditions shifting and leaves senescing, isolates with a fast-growth strategy are able to grow rapidly, dominating the phyllosphere community and leading to its further homogeneization before leaves fully senesce. This scenario provides an explanation for the observation of community convergence and increasing homogeneity of phyllosphere communities throughout the growing season (65, 66).

Our study illustrates that *Methylobacterium* is a complex group of divergent lineages with different ecological strategies and distributions, reflecting long-term adaptation to contrasting local environments. Based upon a similar observation, some authors recently proposed to reclassify *Methylobacterium* group B within a new genus (*Methylorubrum*) that they argue is ecologically and evolutionarily distinct from other *Methylobacterium* clades (31). Although clade B was well supported as a distinct clade in our analyses, our results suggest that it is in fact embedded within clade A, which would render the genus *Methylobacterium* paraphyletic if clade B is defined as a distinct genus (**Figure S1**). Furthermore, group B was not particularly ecologically distinct in comparison with other major clades (**Figure 2**). Our results emphasize the fact that thorough genomic investigations are needed to clarify the taxomonic status of *Methylobacterium*. Beyond any taxonomic considerations, neither clade identity assessed by individual genetic markers nor the tremendous ecological diversity among *Methylobacterium* clades can predict all of the spatial and temporal variation in *Methylobacterium* diversity in nature. In order to define the niches of *Methylobacterium* clades and to understand the metabolic mechanisms underlying their contrasting life strategies, future characterization of their functions and genome structure will be required using phylogenomic approaches.

In conclusion, we find that *Methylobacterium* adaptive responses to local environmental variation in the phyllosphere are driven by both long-term inherited ecological strategies that differ among major clades within the genus, as well by seasonal changes affecting habitat characteristics and community structure in the phyllosphere habitat. Overall, our study combining culture-free and culture-based approaches provides novel insights into the factors driving fine-scale adaptation of microbes to their habitats. In the case of *Methylobacterium*, our approach revealed the particular importance of considering organismal life-history strategies to help understand the fine-scale diversity and dynamics of this ecologically important taxon.

## Acknowledgments

FRQNT (funding), NSERC (funding), Canada Research Chairs (funding), NSF (DEB-1831838), Geneviève Lajoie, Dominique Tardif, Ariane Lafrenière, Hélène Dion-Phénix, Yves Terrat, Kenta Araya (help for sampling), Sylvain Dallaire (help for phenotyping screen), Sergey Stolyar (discussions), Gault Natural Reserve (McGill University), Station Biologique des Laurentides (UdeM), Centre d’Étude de la Forêt (CEF), QCBS and two anonymous reviewers for their helpful suggestions.

## Author contributions

J.B.L., E.S.L., D.C.M., G.B., B.J.S. and S.W.K. planned fieldwork and the experiments. J.B.L. E.S.L., D.C.M. and G.B. performed experiments. J.B.L., E.S.L., S.W.K., D.S. and J.M.S performed bioinformatic analyses. J.A.F., C.J.M. and J.M.S. provided discussion in the early stages of this study. J.B.L. B.J.S. and S.W.K. drafted the manuscript with contributions from C.J.M.

## Data accessibility

raw reads for *16S rRNA* and *rpoB* barcoding on phyllosphere communities (BioProject PRJNA729807; BioSamples SAMN19164946-SAMN19165146) were deposited in NCBI under SRA Accession Numbers SRR14532212-SRR14532451. Partial nucleotide sequences from marker genes obtained by SANGER sequencing on *Methylobacterium* isolates (BioProject PRJNA730554; Biosamples SAMN19190155-SAMN19190401) were deposited in NCBI under GenBank Accession Numbers MZ268514-MZ268593 (*16S rRNA* gene), MZ330152-MZ330358 (*rpoB* gene) and MZ330130-MZ330151 (*sucA*). Bioprojects, Biosamples, SRA and GenBank accession numbers are listed in **Dataset S1**. R codes and related data were deposited on Github (https://github.com/JBLED/methylo-phyllo-diversity).

## Supplemental material legends

**Supplementary Materials and Methods S1** – Detailled materials and methods and supplementary references.

**Figure S1** - ML phylogenetic trees from *sucA* (a) and *rpoB* (b) concatenated hypervariable (HV) regions and consensus clade tree (c).

**Figure S2** - Experimental design of *Methylobacterium* monitoring for growth performance under four temperature treatments.

**Figure S3** - Example of image analysis of *Methylobacterium* monitoring for growth performance under four temperature treatments.

**Figure S4** - Prediction of log normal growth curve, growth rate and yield for 79 isolates incubated under four temperature treatments.

**Figure S5** – *Alphaproteobacteria* (a) and *Methylobacterium* (b) diversity assessed by *rpoB* barcoding, comparison of *Methylobacterium* diversity assessement from *rpoB* barcoding and *16s rRNA* barcoding (c) and comparison of *Methylobacterium* diversity assessment from *rpoB* barcoding and isolation (d).

**Table S1** - Primers used to amplify hyper variable regions in genes *sucA* and *rpoB* and sequence amplification success in 20 *Methylobacterium* isolates from a pilot survey in MSH in august 2017.

**Table S2** - *Methylobacterium* diversity assessed by culture-dependant (isolates; *rpoB* sanger sequencing) and culture-free approaches (*16S rRNA* and *rpoB* barcoding) and comparison between different methods.

**Dataset S1** – a) List of reference *Methylobacteriaceae* genomes used in this study; b) List of phyllosphere samples and their deposited accession numbers; c) List of 80 methylotrophic isolates from MSH (pilot survey in august 2017); d) Clade assignment of 76 *Methylobacterium* isolates from the 2017 pilot survey based on BLAST; e) List of isolates and nucleotide sequence obtained from pilot and timeline surveys, and they deposited accession numbers; f) List of 167 *Methylobacterium* isolates from timeline survey (2018); g) List of *16s rRNA* and *rpoB* barcoding libraries and they deposited accession numbers; h) List of *16s rRNA* ASV nucleotide sequences, taxonomy and their absolute abundance in 46 phyllosphere samples; i) Summary of *16s rRNA* ASV taxonomic assignation; j) Tests for phylogenetic association of traits with culture-based estimation of *Methylobacterium* diversity for different phylogenetic depth; k) List of *rpoB* ASV nucleotide sequences, taxonomy and their absolute abundance in 184 phyllosphere samples; l) Summary of *rpoB* ASV taxonomic assignation; m) Detail of ANOVA analysis for each *Methobacterium* ASV relative abundance; n) Average rate and yield values for 79 *Methylobacterium* isolates monitored under four temperature treatments; o) Detailled ANOVA results for yield and growth rate.

## References

1. Vorholt JA. 2012. Microbial life in the phyllosphere. 12. Nature Reviews Microbiology 10:828–840.

2. Lindow SE, Brandl MT. 2003. Microbiology of the Phyllosphere. Appl Environ Microbiol 69:1875–1883.

3. Fürnkranz M, Wanek W, Richter A, Abell G, Rasche F, Sessitsch A. 2008. Nitrogen fixation by phyllosphere bacteria associated with higher plants and their colonizing epiphytes of a tropical lowland rainforest of Costa Rica. 5. The ISME Journal 2:561–570.

4. Laforest-Lapointe I, Messier C, Kembel SW. 2016. Host species identity, site and time drive temperate tree phyllosphere bacterial community structure. Microbiome 4:27.

5. Laforest-Lapointe I, Messier C, Kembel SW. 2016. Tree phyllosphere bacterial communities: exploring the magnitude of intra- and inter-individual variation among host species. PeerJ 4:e2367.

6. Noble AS, Noe S, Clearwater MJ, Lee CK. 2020. A core phyllosphere microbiome exists across distant populations of a tree species indigenous to New Zealand. PLOS ONE 15:e0237079.

7. Kembel SW, O’Connor TK, Arnold HK, Hubbell SP, Wright SJ, Green JL. 2014. Relationships between phyllosphere bacterial communities and plant functional traits in a neotropical forest. PNAS 111:13715–13720.

8. Shade A, Gregory Caporaso J, Handelsman J, Knight R, Fierer N. 2013. A meta-analysis of changes in bacterial and archaeal communities with time. 8. The ISME Journal 7:1493–1506.

9. Malik AA, Martiny JBH, Brodie EL, Martiny AC, Treseder KK, Allison SD. 2020. Defining trait-based microbial strategies with consequences for soil carbon cycling under climate change. 1. The ISME Journal 14:1–9.

10. Moyes AB, Kueppers LM, Pett□Ridge J, Carper DL, Vandehey N, O’Neil J, Frank AC. 2016. Evidence for foliar endophytic nitrogen fixation in a widely distributed subalpine conifer. New Phytologist 210:657–668.

11. Lajoie G, Kembel SW. 2019. Making the Most of Trait-Based Approaches for Microbial Ecology. Trends in Microbiology 27:814–823.

12. Nemergut DR, Schmidt SK, Fukami T, O’Neill SP, Bilinski TM, Stanish LF, Knelman JE, Darcy JL, Lynch RC, Wickey P, Ferrenberg S. 2013. Patterns and Processes of Microbial Community Assembly. Microbiol Mol Biol Rev 77:342–356.

13. Cavender□Bares J, Kozak KH, Fine PVA, Kembel SW. 2009. The merging of community ecology and phylogenetic biology. Ecology Letters 12:693–715.

14. Martiny JBH, Jones SE, Lennon JT, Martiny AC. 2015. Microbiomes in light of traits: A phylogenetic perspective. Science 350.

15. Tromas N, Taranu ZE, Castelli M, Pimentel JSM, Pereira DA, Marcoz R, Shapiro BJ, Giani A. 2020. The evolution of realized niches within freshwater Synechococcus. Environmental Microbiology 22:1238–1250.

16. Tromas N, Taranu ZE, Martin BD, Willis A, Fortin N, Greer CW, Shapiro BJ. 2018. Niche Separation Increases With Genetic Distance Among Bloom-Forming Cyanobacteria. Front Microbiol 9.

17. Thompson LR, Sanders JG, McDonald D, Amir A, Ladau J, Locey KJ, Prill RJ, Tripathi A, Gibbons SM, Ackermann G, Navas-Molina JA, Janssen S, Kopylova E, Vázquez-Baeza Y, González A, Morton JT, Mirarab S, Zech Xu Z, Jiang L, Haroon MF, Kanbar J, Zhu Q, Jin Song S, Kosciolek T, Bokulich NA, Lefler J, Brislawn CJ, Humphrey G, Owens SM, Hampton-Marcell J, Berg-Lyons D, McKenzie V, Fierer N, Fuhrman JA, Clauset A, Stevens RL, Shade A, Pollard KS, Goodwin KD, Jansson JK, Gilbert JA, Knight R. 2017. A communal catalogue reveals Earth’s multiscale microbial diversity. 7681. Nature 551:457–463.

18. Poretsky R, Rodriguez-R LM, Luo C, Tsementzi D, Konstantinidis KT. 2014. Strengths and Limitations of 16S rRNA Gene Amplicon Sequencing in Revealing Temporal Microbial Community Dynamics. PLOS ONE 9:e93827.

19. Ranjan R, Rani A, Metwally A, McGee HS, Perkins DL. 2016. Analysis of the microbiome: Advantages of whole genome shotgun versus 16S amplicon sequencing. Biochemical and Biophysical Research Communications 469:967–977.

20. Corpe WA, Rheem S. 1989. Ecology of the methylotrophic bacteria on living leaf surfaces. FEMS Microbiol Ecol 5:243–249.

21. Keppler F, Hamilton JTG, Braß M, Röckmann T. 2006. Methane emissions from terrestrial plants under aerobic conditions. 7073. Nature 439:187–191.

22. Knief C, Ramette A, Frances L, Alonso-Blanco C, Vorholt JA. 2010. Site and plant species are important determinants of the Methylobacterium community composition in the plant phyllosphere. ISME J 4:719–728.

23. Clarke PH. 1983. The biochemistry of Methylotrophs by C. Anthony Academic Press; London, New York, 1982 xvi + 432 pages. £24.00, $49.50. FEBS Letters 160:303–303.

24. Anthony C. 1991. Assimilation of carbon by methylotrophs. Biotechnology 18:79–109.

25. Ivanova EG, Doronina NV, Trotsenko YuA. 2001. Aerobic Methylobacteria Are Capable of Synthesizing Auxins. Microbiology 70:392–397.

26. Madhaiyan M, Poonguzhali S, Lee HS, Hari K, Sundaram SP, Sa TM. 2005. Pink-pigmented facultative methylotrophic bacteria accelerate germination, growth and yield of sugarcane clone Co86032 (Saccharum officinarum L.). Biol Fertil Soils 41:350–358.

27. Madhaiyan M, Poonguzhali S, Sa T. 2007. Metal tolerating methylotrophic bacteria reduces nickel and cadmium toxicity and promotes plant growth of tomato (Lycopersicon esculentum L.). Chemosphere 69:220–228.

28. Dourado MN, Aparecida Camargo Neves A, Santos DS, Araújo WL. 2015. Biotechnological and Agronomic Potential of Endophytic Pink-Pigmented Methylotrophic Methylobacterium spp. Biomed Res Int 2015.

29. Ryu JH (Chungbuk NU, Madhaiyan M (Chungbuk NU, Poonguzhali S (Chungbuk NU, Yim WJ (Chungbuk NU, Indiragandhi P (Chungbuk NU, Kim KA (Chungbuk NU, Anandham R (Chungbuk NU, Yun JC (National I of AS and T, Kim KH (The U of S, Sa TM (Chungbuk NU. 2006. Plant Growth Substances Produced by Methylobacterium spp. and Their Effect on Tomato (Lycopersicon esculentum L.) and Red Pepper (Capsicum annuum L.) Growth. Journal of Microbiology and Biotechnology.

30. Lee HS, Madhaiyan M, Kim CW, Choi SJ, Chung KY, Sa TM. 2006. Physiological enhancement of early growth of rice seedlings (Oryza sativa L.) by production of phytohormone of N2-fixing methylotrophic isolates. Biol Fertil Soils 42:402–408.

31. Green PN, Ardley JK. 2018. Review of the genus Methylobacterium and closely related organisms: a proposal that some Methylobacterium species be reclassified into a new genus, Methylorubrum gen. nov. International Journal of Systematic and Evolutionary Microbiology 68:2727–2748.

32. Chen W-M, Cai C-Y, Li Z-H, Young C-C, Sheu S-Y. 2019. Methylobacterium oryzihabitans sp. nov., isolated from water sampled from a rice paddy field. International Journal of Systematic and Evolutionary Microbiology, 69:3843–3850.

33. Feng G-D, Chen W, Zhang X-J, Zhang J, Wang S-N, Zhu H. 2020. Methylobacterium nonmethylotrophicum sp. nov., isolated from tungsten mine tailing. International Journal of Systematic and Evolutionary Microbiology, 70:2867–2872.

34. Jia L juan, Zhang K shuai, Tang K, Meng J yu, Zheng C, Feng F ying. 2020. Methylobacterium crusticola sp. nov., isolated from biological soil crusts. International Journal of Systematic and Evolutionary Microbiology, 70:2089–2095.

35. Kim J, Chhetri G, Kim I, Lee B, Jang W, Kim MK, Seo T. 2020. Methylobacterium terricola sp. nov., a gamma radiation-resistant bacterium isolated from gamma ray-irradiated soil. International Journal of Systematic and Evolutionary Microbiology, 70:2449–2456.

36. Kim J, Chhetri G, Kim I, Kim MK, Seo T. 2020. Methylobacterium durans sp. nov., a radiation-resistant bacterium isolated from gamma ray-irradiated soil. Antonie van Leeuwenhoek 113:211–220.

37. Ten LN, Li W, Elderiny NS, Kim MK, Lee S-Y, Rooney AP, Jung H-Y. 2020. Methylobacterium segetis sp. nov., a novel member of the family Methylobacteriaceae isolated from soil on Jeju Island. Arch Microbiol 202:747–754.

38. Jiang L, An D, Wang X, Zhang K, Li G, Lang L, Wang L, Jiang C, Jiang Y. 2020. Methylobacterium planium sp. nov., isolated from a lichen sample. Arch Microbiol 202:1709–1715.

39. Pascual JA, Ros M, Martínez J, Carmona F, Bernabé A, Torres R, Lucena T, Aznar R, Arahal DR, Fernández F. 2020. Methylobacterium symbioticum sp. nov., a new species isolated from spores of Glomus iranicum var. tenuihypharum. Curr Microbiol 77:2031–2041.

40. Marx CJ, Bringel F, Chistoserdova L, Moulin L, Farhan Ul Haque M, Fleischman DE, Gruffaz C, Jourand P, Knief C, Lee M-C, Muller EEL, Nadalig T, Peyraud R, Roselli S, Russ L, Goodwin LA, Ivanova N, Kyrpides N, Lajus A, Land ML, Médigue C, Mikhailova N, Nolan M, Woyke T, Stolyar S, Vorholt JA, Vuilleumier S. 2012. Complete Genome Sequences of Six Strains of the Genus Methylobacterium. J Bacteriol 194:4746–4748.

41. Tani A, Ogura Y, Hayashi T, Kimbara K. 2015. Complete Genome Sequence of Methylobacterium aquaticum Strain 22A, Isolated from Racomitrium japonicum Moss. Genome Announc 3.

42. Minami T, Ohtsubo Y, Anda M, Nagata Y, Tsuda M, Mitsui H, Sugawara M, Minamisawa K. 2016. Complete Genome Sequence of Methylobacterium sp. Strain AMS5, an Isolate from a Soybean Stem. Genome Announc 4.

43. Morohoshi T, Ikeda T. 2016. Complete Genome Sequence of Methylobacterium populi P-1M, Isolated from Pink-Pigmented Household Biofilm. Genome Announc 4.

44. Belkhelfa S, Labadie K, Cruaud C, Aury J-M, Roche D, Bouzon M, Salanoubat M, Döring V. 2018. Complete Genome Sequence of the Facultative Methylotroph Methylobacterium extorquens TK 0001 Isolated from Soil in Poland. Genome Announc 6.

45. Green PN. 2015. Methylobacterium, p. 1–8. In Bergey’s Manual of Systematics of Archaea and Bacteria. American Cancer Society.

46. Charron G, Leducq JB, Bertin C, Dube AK, Landry CR. 2014. Exploring the northern limit of the distribution of Saccharomyces cerevisiae and Saccharomyces paradoxus in North America. FEMS Yeast Res 14:281–8.

47. Vos M, Quince C, Pijl AS, Hollander M de, Kowalchuk GA. 2012. A Comparison of rpoB and 16S rRNA as Markers in Pyrosequencing Studies of Bacterial Diversity. PLOS ONE 7:e30600.

48. Ogier J-C, Pagès S, Galan M, Barret M, Gaudriault S. 2019. rpoB, a promising marker for analyzing the diversity of bacterial communities by amplicon sequencing. BMC Microbiol 19:171.

49. Ronquist F, Teslenko M, van der Mark P, Ayres DL, Darling A, Höhna S, Larget B, Liu L, Suchard MA, Huelsenbeck JP. 2012. MrBayes 3.2: Efficient Bayesian Phylogenetic Inference and Model Choice Across a Large Model Space. Systematic Biology 61:539–542.

50. Küpfer M, Kuhnert P, Korczak BM, Peduzzi R, Demarta A. 2006. Genetic relationships of Aeromonas strains inferred from 16S rRNA, gyrB and rpoB gene sequences. International Journal of Systematic and Evolutionary Microbiology, 56:2743–2751.

51. Ee R, Madhaiyan M, Ji L, Lim Y-L, Nor NM, Tee K-K, Chen J-W, Yin W-F. 2016. Chania multitudinisentens gen. nov., sp. nov., an N-acyl-homoserine-lactone-producing bacterium in the family Enterobacteriaceae isolated from landfill site soil. International Journal of Systematic and Evolutionary Microbiology, 66:2297–2304.

52. Frank RAW, Price AJ, Northrop FD, Perham RN, Luisi BF. 2007. Crystal Structure of the E1 Component of the Escherichia coli 2-Oxoglutarate Dehydrogenase Multienzyme Complex. Journal of Molecular Biology 368:639–651.

53. Bell RL, González-Escalona N, Stones R, Brown EW. 2011. Phylogenetic evaluation of the ‘Typhimurium’ complex of Salmonella strains using a seven-gene multi-locus sequence analysis. Infection, Genetics and Evolution 11:83–91.

54. Dixon P. 2003. VEGAN, a package of R functions for community ecology. Journal of Vegetation Science 14:927–930.

55. Redford AJ, Bowers RM, Knight R, Linhart Y, Fierer N. 2010. The ecology of the phyllosphere: geographic and phylogenetic variability in the distribution of bacteria on tree leaves. Environmental Microbiology 12:2885–2893.

56. Callahan BJ, McMurdie PJ, Rosen MJ, Han AW, Johnson AJA, Holmes SP. 2016. DADA2: High resolution sample inference from Illumina amplicon data. Nat Methods 13:581–583.

57. Quast C, Pruesse E, Yilmaz P, Gerken J, Schweer T, Yarza P, Peplies J, Glöckner FO. 2013. The SILVA ribosomal RNA gene database project: improved data processing and web-based tools. Nucleic Acids Res 41:D590–D596.

58. Boratyn GM, Camacho C, Cooper PS, Coulouris G, Fong A, Ma N, Madden TL, Matten WT, McGinnis SD, Merezhuk Y, Raytselis Y, Sayers EW, Tao T, Ye J, Zaretskaya I. 2013. BLAST: a more efficient report with usability improvements. Nucleic Acids Res 41:W29–33.

59. Stegen JC, Lin X, Fredrickson JK, Chen X, Kennedy DW, Murray CJ, Rockhold ML, Konopka A. 2013. Quantifying community assembly processes and identifying features that impose them. ISME J 7:2069–2079.

60. Kembel SW, Cowan PD, Helmus MR, Cornwell WK, Morlon H, Ackerly DD, Blomberg SP, Webb CO. 2010. Picante: R tools for integrating phylogenies and ecology. Bioinformatics 26:1463–1464.

61. Kembel SW. 2009. Disentangling niche and neutral influences on community assembly: assessing the performance of community phylogenetic structure tests. Ecology Letters 12:949–960.

62. Zwietering MH, Jongenburger I, Rombouts FM, Riet K van ‘t. 1990. Modeling of the Bacterial Growth Curve. Appl Environ Microbiol 56:1875–1881.

63. Lipson DA. 2015. The complex relationship between microbial growth rate and yield and its implications for ecosystem processes. Front Microbiol 6.

64. Cavender Bares J, Ackerly DD, Baum DA, Bazzaz FA. 2004. Phylogenetic Overdispersion in Floridian Oak Communities. The American Naturalist 163:823–843.

65. Copeland JK, Yuan L, Layeghifard M, Wang PW, Guttman DS. 2015. Seasonal Community Succession of the Phyllosphere Microbiome. MPMI 28:274–285.

66. Maignien L, DeForce EA, Chafee ME, Eren AM, Simmons SL. Ecological Succession and Stochastic Variation in the Assembly of Arabidopsis thaliana Phyllosphere Communities. mBio 5:e00682–13.

