## Supplementary material for "Fine-Scale Adaptations to Environmental Variation and Growth Strategies Drive Phyllosphere *Methylobacterium* Diversity": Table S1

**Table S1 - Primers used to amplify hyper variable regions in genes *sucA* and *rpoB.*** Sequence amplification success (%) in 20 *Methylobacterium* isolates from a pilot survey in MSH in august 2017 (see **Dataset S1c).** These genes were used as an alternative to the *16S rRNA* gene to develop a highly polymorphic marker targeting specifically the *Methylobacteriaceae* family. For each of the five hyper variable regions (three in *sucA*, two in *rpoB*), primers were designed in flanking by well-conserved regions.

| **Clade** | | | **A1** | **A2** | **A6** | **A9** | **A10** |
| --- | --- | --- | --- | --- | --- | --- | --- |
| **Isolates** | | | DNA001 DNA006 DNA010 DNA014 DNA018 DNA024 LYS027 LYS051 LYS083 | LYS069 LYS093 | DNA007 DNA013 LYS037 | DNA011 DNA012 LYS072 LYS080 | DNA020 DNA021 |
| ***sucA*** | Met01-3-F GCGCAGGATGTGGAAGTAG | Met01-960-R SATCGACATGCTSTGCTACC | 89% | 100% | 100% | 50% | 100% |
|  | Met01-306-F YTCSGAGAGCATCGAGTTG | Met01-1160-R GGMTCGRTCCACTTCATCA | 100% | 50% | 100% | 25% | 100% |
|  | Met01-1035-F GCCGTTGCAGTGGAAGAT | Met01-1758-R GGGCAAGGACAAGGARATY | 89% | 100% | 100% | 100% | 100% |
| ***rpoB*** | **Met02-352-F***  **AAGGACATCAAGGAGCAGGA** | **Met02-1121-R***  **ACSCGGTAKATGTCGAACAG** | **100%** | **100%** | **100%** | **100%** | **100%** |
|  | Met02-2480-F  TGAAGGATGACGTGTTCACC | Met02-3116-R  TTCGACTCGTCGTACTGCTT | 100% | 100% | 100% | 100% | 100% |
| *** Combination of primers selected as marker for the rest of this study** | | | | | | | |
