## Supplementary material for "Fine-Scale Adaptations to Environmental Variation and Growth Strategies Drive Phyllosphere *Methylobacterium* Diversity": Table S2

**Table S2 - *Methylobacterium* diversity assessed by culture-dependant (isolates; *rpoB* sanger sequencing) and culture-free approaches (*16S rRNA* and *rpoB* barcoding) and comparison between different methods**. For each clade (unassigned marked as “un”), the following information is shown: number of ASVs and their relative abundance (F) estimated by *16* *rRNA* barcoding in 46 phyllosphere samples (**Dataset S1i**); the number of isolates (*rpoB* SANGER sequencing) obtained from 76 phyllosphere samples (**Dataset S1f**); the number of ASVs and their relative abundance estimated by *rpoB* barcoding in 184 phyllosphere samples (**Dataset S1k,l**); the comparison of diversities assessed by *rpoB* barcoding and isolates assuming either 100% or 98.5% *rpoB* nucleotide sequence identity between isolates and ASVs. In each case, the relative abundance F was calculated after removing non-*Methylobacterium* sequences.

| **Clade** | ***16S rRNA***  **ASVs** | ***16S rRNA***  **ASVs abundance (F)** | **Isolates** | ***rpoB***  **ASVs** | ***rpoB* ASV abundance (F)** | **100% *rpoB***  **sequence match** | | | **98.5% *rpoB***  **sequence match** | | |
| --- | --- | --- | --- | --- | --- | --- | --- | --- | --- | --- | --- |
|  |  |  |  |  |  | ***rpoB***  **ASVs** | **Isolates** | **F** | ***rpoB***  **ASVs** | **Isolates** | **F** |
| **A1** | 3 | 0.097 | 9 | 31 | 0.066 | 4 | 4 | 0.022 | 26 | 7 | 0.063 |
| **A2** | - | - | 3 | 7 | 0.003 | 1 | 1 | 0.001 | 4 | 1 | 0.002 |
| **A3** | - | - | - | 2 | 0.001 | - | - | - | - | - | - |
| **A4** | - | - | - | 20 | 0.017 | - | - | - | - | - | - |
| **A5** | - | - | 1 | 3 | 0.001 | 1 | 1 | 0.001 | 1 | 1 | 0.001 |
| **A6** | 1 | 0.224 | 41 | 37 | 0.243 | 16 | 28 | 0.192 | 27 | 40 | 0.232 |
| **A9** | 9 | 0.676 | 100 | 59 | 0.452 | 27 | 83 | 0.387 | 54 | 95 | 0.434 |
| **A10** | 2 | 0.003 | 6 | 6 | 0.010 | 2 | 3 | 0.006 | 2 | 4 | 0.006 |
| **B** | - | - | 7 | 31 | 0.191 | 2 | 3 | 0.105 | 10 | 7 | 0.122 |
| **un** | - | - | - | 4 | 0.015 | - | - | - | - | - | - |
| **Total** | 15 | 1 | 167 | 200 | 1 | 53 | 123 | 0.712 | 124 | 155 | 0.859 |
| **Part of total diversity match in *rpoB* comparisons** | | | | | | **26.5%** | **73.7%** | **71.2%** | **62.0%** | **92.8%** | **85.9%** |
