## Supplementary material for "Fine-Scale Adaptations to Environmental Variation and Growth Strategies Drive Phyllosphere *Methylobacterium* Diversity": Figure S1

**Figure S1 - ML phylogenetic trees from *sucA* (a) and *rpoB* (b) concatenated hypervariable (HV) regions.** Trees were drawn from sequences obtained for 20 representative isolates from 2017 pilot survey (black circles) and reference genomes. Trees with the highest log likelihood are shown. Bootstraps: only values for node supported by at least 50% of replicated trees are displayed. Phylogenetic tree was rooted on *Microvirga* and *Enterovirga* outgroups (Compressed). a) The *sucA* ML tree was inferred from 3 aligned concatenated HV regions (1,663 bp) available for 189 reference genomes and 14 tested isolates. b) The *rpoB* ML tree was inferred from 2 aligned and concatenated HV regions (1,244 bp) available for 163 reference genomes and the 20 tested isolates. c) Consensus clade tree from *sucA* and *rpoB* ML phylogenies. Only tree topology among clades supported by both phylogenies is shown, regardless bootstrap support. For each consensus node, the minimum (most conservative) bootstrap support found between phylogenies is shown (grey scale, legend on top).

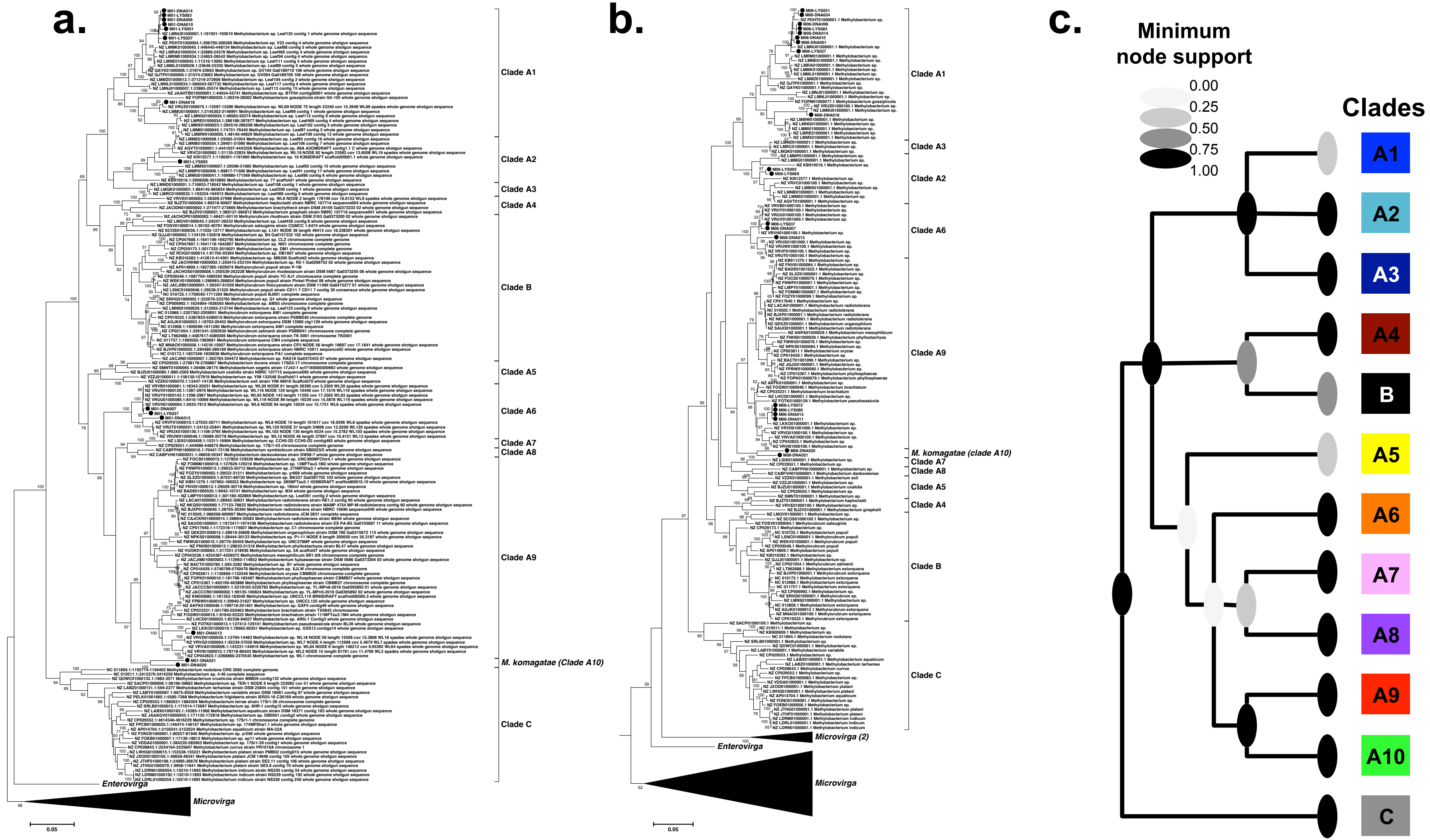
a.
Clade A1
Clade A2
Clade A3
Clade A4
Clade B
Clade A5
Clade A6
Clade A7
Clade A8
Clade A9
M. komagatae (Clade A10)
Clade C
Enterovirga
Microvirga
0.05
b.
Clade A1
Clade A3
Clade A2
Clade A6
Clade A9
M. komagatae (clade A10)
Clade A7
Clade A8
Clade A5
Clade A4
Clade B
Clade C
Microvirga (2)
Enterovirga
Microvirga
0.05
c.
Minimum node support
0.00
0.25
0.50
0.75
1.00
Clades
A1
A2
A3
A4
B
A5
A6
A7
A8
A9
A10
C
