## Supplementary material for "Fine-Scale Adaptations to Environmental Variation and Growth Strategies Drive Phyllosphere *Methylobacterium* Diversity": Figure S2

**Figure S2 - Experimental design of *Methylobacterium* monitoring for growth performance under four temperature treatments**. a) 79 isolates (two showed in this example: pink and cyan) and two negative controls (not showed) were tested for ability to grow under different temperature treatments. b) Pre-conditioning step (*P*): For each isolate and negative controls and each temperature treatment (20 and 30 °C), 10µL of cellular culture from stock were spread on solid Methanol-MMS media. c) After 20 days of incubation at 20 °C (*P20*) or 30 °C (*P30*), petri dishes were swabbed and collected cell concentrations adjusted to $OD_{630}$=0.2. d) Monitoring step (*M*): Each pre-conditioned culture *P20* (*n*=81) and *P30* (*n*=81), was spotted on new Methanol-MMS media (five replicates per culture, per *P* treatment). Each petri dish was duplicated, one copy for incubation at 20 °C (*M20* treatment) and one for incubation at 30 °C (*M30*). Pictures of petri dishes were took 7, 13 and 24 after inoculation. e) Three examples of spot organization on petri dishes (24 per *M* treatment, 17 isolates + one negative control per dish). Open circle represent negative controls.

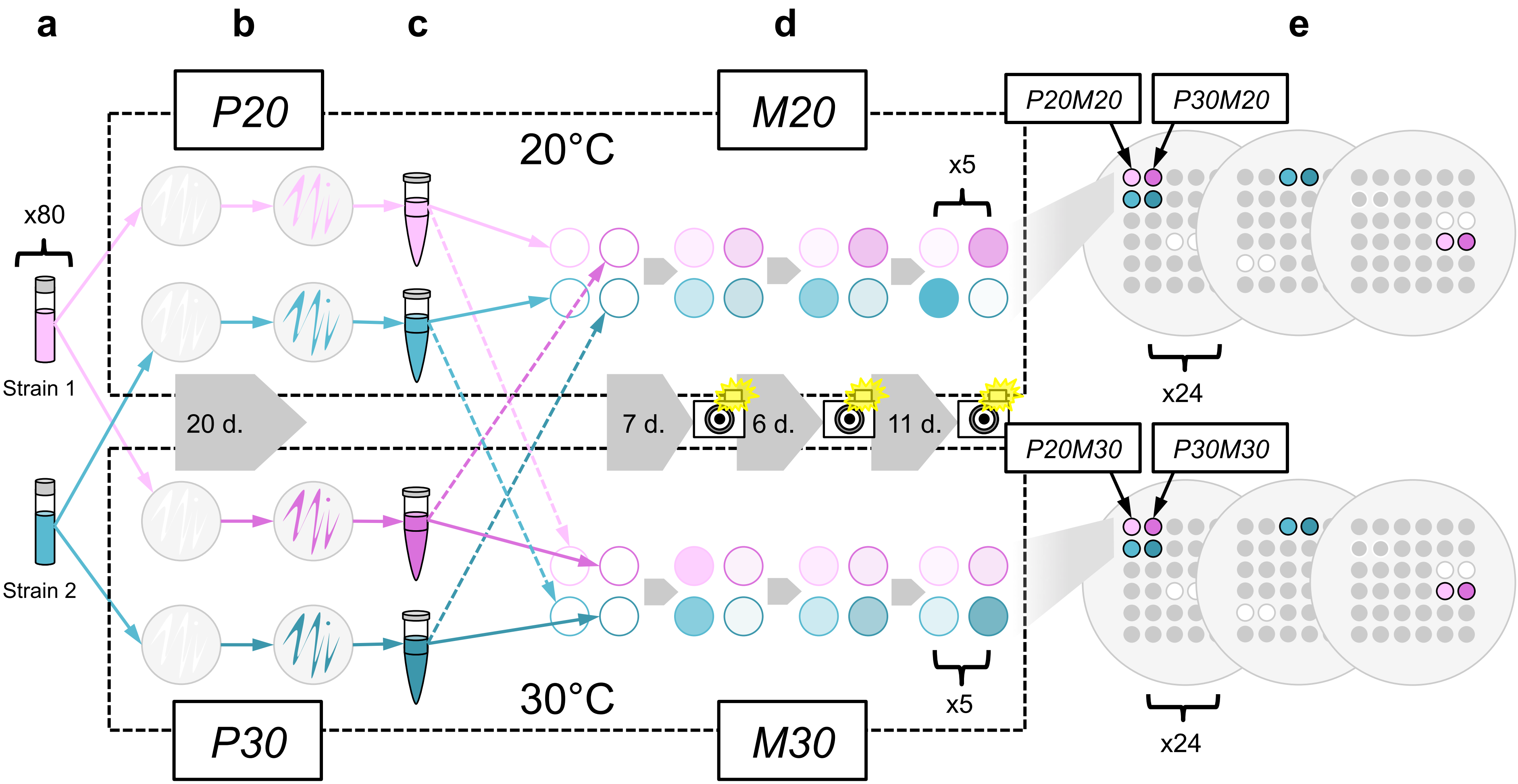

a
b
c
d
e
P20
M20
P20M20
P30M20
20°C
x80
x5
Strain 1
20 d.
7 d.
6 d.
11 d.
x24
P20M30
P30M30
Strain 2
30°C
x5
x24
P30
M30
