## Supplementary material for "Fine-Scale Adaptations to Environmental Variation and Growth Strategies Drive Phyllosphere *Methylobacterium* Diversity": Figure S3

**Figure S3 - Example of image analysis of *Methylobacterium* monitoring for growth performance under four temperature treatments**. a) Original picture. b) The original picture was converted in grey scale in ImajeJ. Areas outside of the agar, as well as every visible particle other than bacteria spot within the agar area, were manually cropped (black). The picture was duplicated. A copy was used for background correction (*BACK*; bacteria spots cropped). Another copy was used to measure raw bacteria spot intensities (*BW*). c) Reconstruction of background intensities. d) Correction of raw intensities by subtracting background values. e) Definition of growth area: detail of a spot (top) and average pixel intensities in function of the radius of concentric circles drawn from the center of the colony. Growth area is defined as the circle with maximal *T* value in *t*-test comparison between intensities outside and within the area (here in red). f) Comparison of intensity distributions outside (red; local background) and within (blue; spot) growing area. Dotted lines indicate average intensity values. g) Border effect: spot intensities after correction for local background (*I*) are shown for 48 petri dishes and 3 time points in function of their average position of each petri. Because of less competition for nutrients, *I* values (proportional to point size) are in average higher close to the border of the petri dish (spot positioned according to the original picture). h) Expected *I* values (scale on top) in function of *X*/*Y* position on the petri dish predicted from a polynomial regression ($I \sim X^2Y^2+X^2Y+XY^2+X+Y$). i) Corrected *I* values (residuals from the polynomial regression).

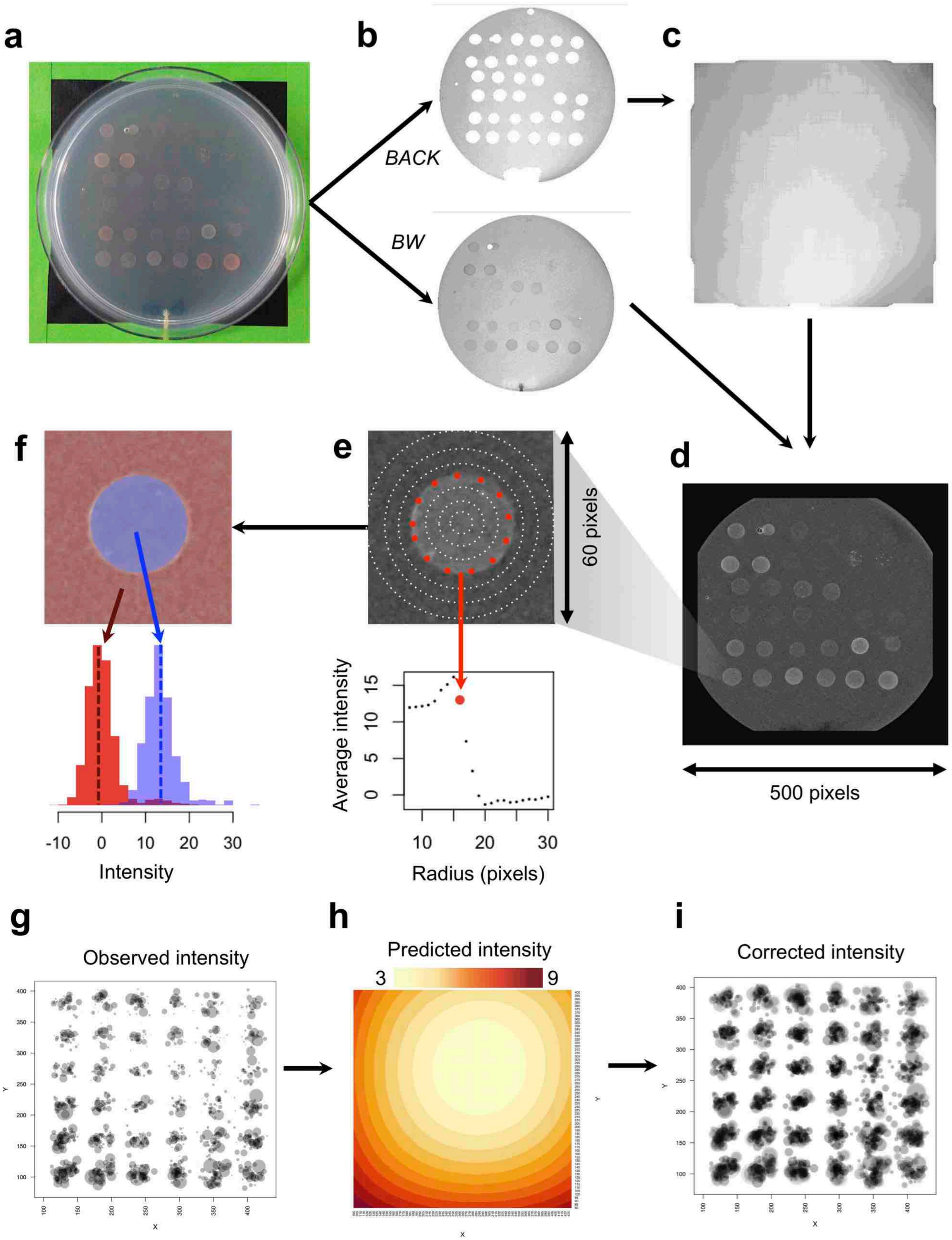
a
b
BACK
BW
c
d
500 pixels
e
60 pixels
Average intensity
Radius (pixels)
f
Intensity
g
Observed intensity
h
Predicted intensity
3
9
i
Corrected intensity
