## Supplementary material for "Fine-Scale Adaptations to Environmental Variation and Growth Strategies Drive Phyllosphere *Methylobacterium* Diversity": Figure S4

**Figure S4 - Prediction of log normal growth curve, growth rate and yield for 79 isolates incubated under four temperature treatments.** a) Log normal best prediction for four different observed cases (legend on bottom). Curves were predicted in the range 0-36 days form values observed at $T_7$, $T_{13}$ and $T_{24}$, assuming null intensity at $T_0$. Models assuming that intensity remained null until $T_i$ were tested in the range 0-7 days. b) determination of yield (*Y*, maximum intensity) and growth rate ($r = 1/log+lag$) from predicted growth curve. c) comparison of predicted yield and maximum observed intensity. d) comparison of predicted log+lag values with time ($T_7$, $T_{13}$ or $T_{24}$) at which maximum intensity was observed. e) All predicted growth curves showed separately for each temperature treatment. Replicates for which maximum intensity was not reach at day 36 according to the model were discarded ($log+lag \geq 36$).

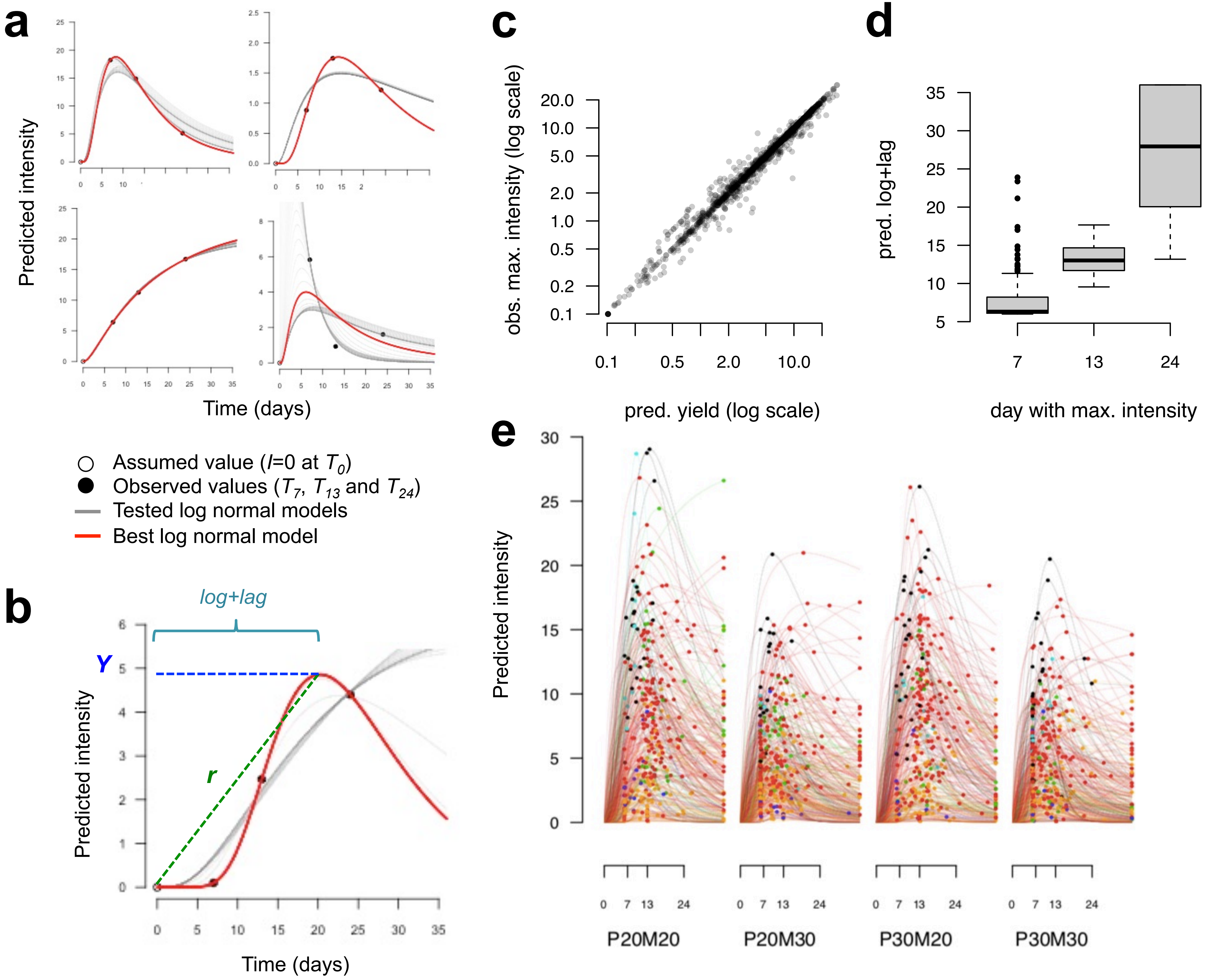
a
Predicted intensity
Time (days)
Assumed value (I=0 at T0)
Observed values (T7, T13 and T24)
Tested log normal models
Best log normal model
b
log+lag
Y
r
Predicted intensity
Time (days)
c
obs. max. intensity (log scale)
pred. yield (log scale)
d
pred. log+lag
day with max. intensity
e
Predicted intensity
P20M20
P20M30
P30M20
P30M30
