## Supplementary material for "Fine-Scale Adaptations to Environmental Variation and Growth Strategies Drive Phyllosphere *Methylobacterium* Diversity": Figure S5

**Figure S5 – *Alphaproteobacteria* and *Methylobacterium* diversity assessed by *rpoB* barcoding.** a) Unrooted ML phylogenetic tree based on *rpoB* partial nucleotide sequences from 1,344 ASVs (410 bp). Only nodes supported by at least 50% of bootstraps (200 permutations) are shown. ASVs are labeled according to their taxonomic assignation based on *rpoB* nucleotide sequence database (legend on bottom left). b) Unrooted ML phylogenetic tree based on *rpoB* partial nucleotide sequences from 283 *Methylobacteriaceae* ASV (points) and 232 references isolates and genomes (unlabeled tips). Only nodes supported by at least 50% of bootstraps (200 permutations) are shown. Full circle indicate 200 *Methylobacterium* ASVs assigned to clades (colored) when clustering with identified reference sequences with at least 50% of support. Unassigned ASVs are indicated in grey. c) Heatmap showing the comparison of *Methylobacterium* diversity assessement from *rpoB* barcoding and *16s rRNA* barcoding. For 41 phyllosphere samples and the METH community (in rows; colors indicating sample origin; legend on bottom left), the relative sequence abundances (color scale on top left) of ASVs assigned to the same clade were combined (in columns). For *rpoB* barcoding, all clades were detected (Unk. indicates unassigned ASVs). For *16s rRNA* barcoding only clades A1, A6, A9 and A10 were detected. d) Comparison of *Methylobacterium* diversity assessment from *rpoB* barcoding (Y-axis) and isolation (X-axis). Number of *Methylobacterium* isolates in function of ASV relative abundance assuming 98.5% of nucleotide identity between *rpoB* sequences obtained by SANGER sequencing in isolates and *rpoB* sequences of ASVs (maximum of 6 nucleotide mismatches). Points are colored according to assignement to clades (legend on top right). The proportion of unmatched diversity (no match between ASV and isolate sequences) is displayed in pie charts for ASVs (top left) and isolates (bottom right).

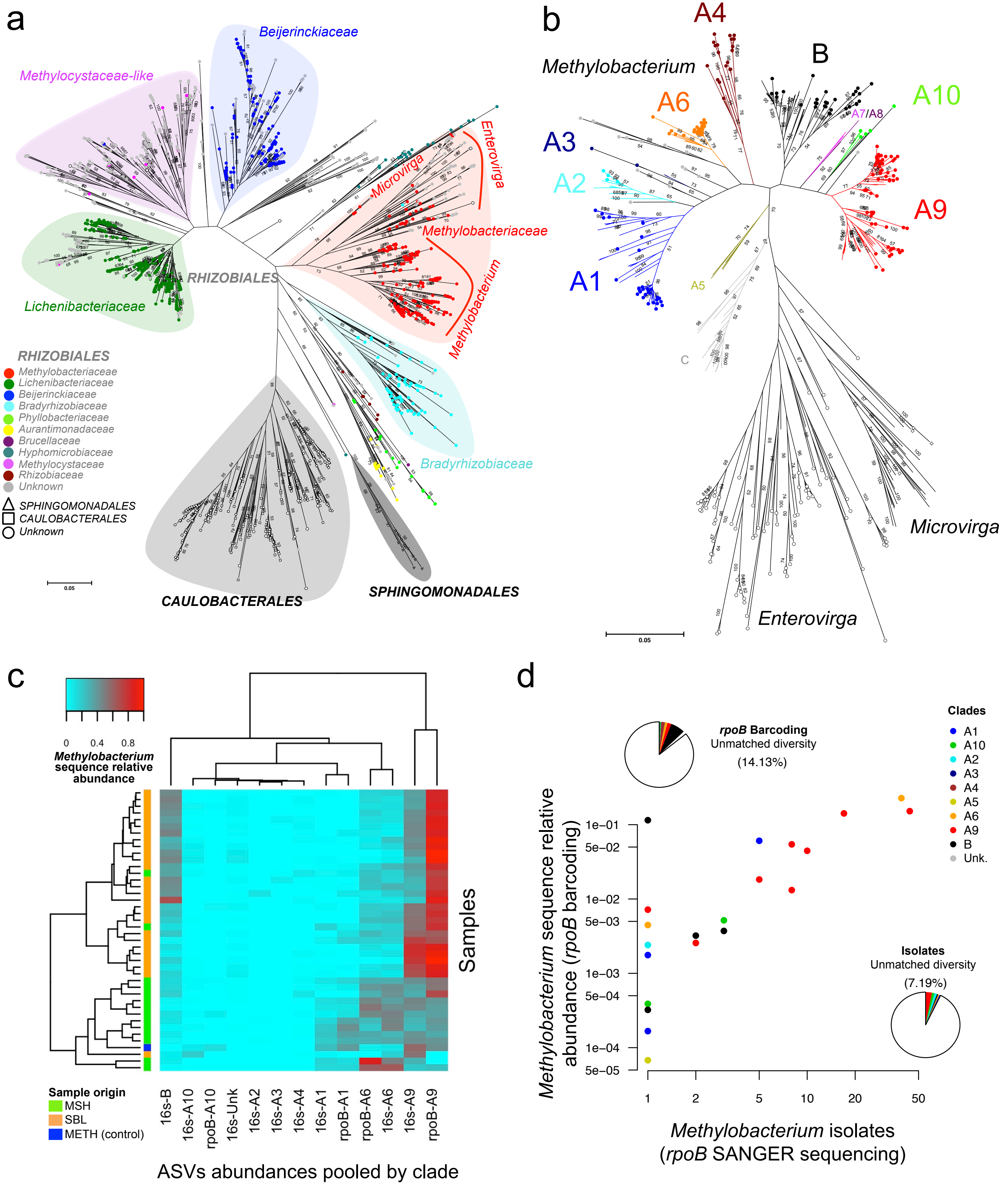

a
Beijerinckiaceae
Methylocystaceae-like
Lichenibacteriaceae
RHIZOBIALES
Microvirga
Enterovirga
Methylobacteriaceae
Methylobacterium
Bradyrhizobiaceae
CAULOBACTERALES
SPHINGOMONADALES
0.05
RHIZOBIALES
Methylobacteriaceae
Lichenibacteriaceae
Beijerinckiaceae
Bradyrhizobiaceae
Phyllobacteriaceae
Aurantimonadaceae
Brucellaceae
Hyphomicrobiaceae
Methylocystaceae
Rhizobiaceae
Unknown
SPHINGOMONADALES
CAULOBACTERALES
Unknown
b
Methylobacterium
A4
B
A6
A10
A7/A8
A3
A2
A9
A1
A5
C
Microvirga
Enterovirga
0.05
c
0
0.4
0.8
Methylobacterium sequence relative abundance
Samples
Sample origin
MSH
SBL
METH (control)
16s-B
16s-A10
rpoB-A10
16s-Unk
16s-A2
16s-A3
16s-A4
16s-A1
rpoB-A1
rpoB-A6
16s-A6
16s-A9
rpoB-A9
ASVs abundances pooled by clade
d
rpoB Barcoding
Unmatched diversity
(14.13%)
Clades
A1
A10
A2
A3
A4
A5
A6
A9
B
Unk.
Isolates
Unmatched diversity
(7.19%)
Methylobacterium sequence relative abundance (rpoB barcoding)
1e−01
5e−02
1e−02
5e−03
1e−03
5e−04
1e−04
5e−05
1
2
5
10
20
50
Methylobacterium isolates (rpoB SANGER sequencing)
