## Supplementary Materials and Methods S1 for "Fine-Scale Adaptations to Environmental Variation and Growth Strategies Drive Phyllosphere *Methylobacterium* Diversity"

#### Phylogenetics of plant-associated *Methylobacterium* diversity

We aimed to assess the known diversity of *Methylobacterium* and its distribution across biomes and especially the phyllosphere. First, we constructed a phylogeny of *Methylobacteriaceae* from the complete sequence of *rpoB*, a highly polymorphic housekeeping gene commonly used to reconstruct robust phylogenies in bacteria, because unlikely to experience horizontal gene transfer or copy number variation (1, 2). We retrieved this gene from all complete and draft *Methylobacteriaceae* genomes publicly available in September 2020, including 153 *Methylobacteria*, 30 *Microvirga* and 2 *Enterovirga* (**Dataset S1a**), using blast of the *rpoB* complete sequence from the *M. extorquens* strain TK001 against NCBI databases *refseq_genomes* and *refseq_rna* (3) available for *Methylobacteriaceae* (Uncultured/environmental samples excluded). Nucleotide sequences were converted in amino-acid in MEGA7 (4, 5), aligned according to the protein sequence, an converted back in nucleotides. The *Methylobacteriaceae* phylogenetic tree was inferred from the *rpoB* nucleotide sequence alignment (4064 bp). Nodal support values (Bayesian posterior probabilities) were estimated using MrBayes v. 3.2.7a (6). Bayesian analyses consisted of paired independent runs, each using four Metropolis coupled chains that consisted of 5 million generations, after which standard deviation of split frequencies had stabilized to less than 0.03. For each of the paired runs, trees were sampled every 1000 generations and the first 1 million generations were treated as the burn-in and discarded. The remaining trees from the two runs (*n*=1802) were combined to determine split frequencies (nodal posterior probabilities). The consensus tree and nodal support values were determined in PAUP v. 4. To improve the presentation of the tree, branch lengths were computed in R, using the Grafen method (7) (function *compute.brlen* in package *ape* ; **Figure 2**). For each *Methylobacterium* reference strain, we retrieved the species name and the sampling origin, when available. Additionally, we assigned each strain to a group (A, B, C) according to previously proposed subdivisions (8). Because group A consisted in several paraphyletic groups branching deeply in the *rpoB* phylogeny, we subdivided *Methylobacterium* group A into 9 clades (A1-A9), using a ~92% pairwise similarity (PS) cut-off on the *rpoB* complete sequence. PS was calculated in MEGA7 from nucleotide sequences as PS = *1-pdistance*.

#### Study sites and phyllosphere sampling

##### Study forests

The two study forests were located in Gault Nature Reserve (Mont Saint-Hilaire, Quebec, Canada ; 45.54 N 73.16 W), here referred as MSH, an old forest occupying the hill of Mount Saint-Hilaire, and *Station Biologique des Laurentides* (Saint-Hippolyte, Quebec, Canada ; 45.99 N 73.99 W), here referred as SBL, a mosaic of natural wetlands, xeric and mesic forests (**Figure 1; Dataset S1b**).

In august 2017, we realized a pilot survey in MSH. We choose two plots (MSH-L0 and MSH-H0) with similar tree species composition and different elevations (175 and 220 m, respectively). Forests were dominated by merican beech (*Fagus grandifolia*), sugar maple (*Acer saccharum*), striped maple (*Acer pensylvanicum*), birch (*Betula alleghaniensis*) and northern red oak (*Quercus rubra*). We collected leaves from 18 randomly chosen trees among dominant species (9 trees per forest), for which we were able to sample the lower part of the canopy (3-5m), hence excluding *B. alleghaniensis* and *Q. rubra*. Additionally, we sampled one American hophornbeam (*Ostrya virginiana*) in MSH-H0. For each tree (n=19), sampling was replicated tree times in different parts of the subcanopy (3-5m), whenever possible.

In 2018, we realized a timeline survey in MSH and SBL. In MSH, we choose 6 plots distributed along a 1.2 km ecological and altitudinal transect (MSH1-6). Lower plots MSH1,6 (170-190 m) were dominated by *F. grandifolia* and *A. saccharum*. Medium plots MSH2,3 (225-270 m) were dominated by *B. alleghaniensis*, *Q. rubra* and *A. saccharum*. Higher plots MSH4,5 (290-315 m) were dominated by *A. saccharum*, *Q. rubra* and *O. virginiana*. In SBL, we choose 4 plots (SBL1-4) distributed along a 1.2 km transect in a transition zone dominated by *A. saccharum*, *F. grandifolia*, *A. pensylvanicum,* balsam fir (*Abies balsamea*), red maple (*Acer rubrum*), and paper birch (*Betula papyrifera*). During this aurvey, we tagged 80 trees (40 per forest, 6-10 per plots) from the dominant species for which we were able to sample the subcanopy, hence excluding *Betula ssp* from any plot and *Q. rubra* from MSH2,3. Each tree was sampled 3-4 times from June to October 2018. For each time point and each tree, sampling was replicated twice in different parts of the subcanopy, whenever possible.

#### Sample collection

Sampling consisted of collecting about 2-10g of leaves into Fisherbrand® sterile bags with a pole pruner while wearing nitrile gloves. For each plot, we realized a negative controls consisting in empty sterile bags opened and sealed on site. Samples were sealed and conserved up to 48h at 4°C until further processing. Samples were randomized before processing. Each bag was unsealed under a sterile hood and 50ml of sterile phosphate buffer was added ($KH_2PO_4$ 100mM solution poured into $K_2HPO_4$ 100mM solution until pH 7.3). Bags were sealed again and vigorously agitated for 5 minutes. For each bag, about 45 ml of phosphate buffer containing the microbial community was transferred in sterile 50 ml Falcon® tubes and centrifuged for 30 minutes at 3,900 rpm at 4°C. Supernatant was removed in order to left the pellet in less than 1 ml of phosphate buffer. The pellet was resuspended and split in two equal volumes in two sterile 1.5 ml tubes. The first tube was directly stored at -80 °C for future DNA extraction, metagenomics and community analysis. The second tube was completed with 500 µl of 50% glycerol (minimum final glycerol concentration: 25%) for future isolations and stored at -80 °C.

Samples were randomized before DNA extraction. DNA extraction was performed using a DNeasy PowerSoil Pro Kit (Qiagen) with the following modifications: each sample (~500 µl) was thawed on ice and centrifuged in order to resuspend the pellet containing the microbial community in a maximum 200 µl volume before proceeding to extraction. DNA was eluted in 50µl of solution C6 and stored at -20 °C. DNA was quantified using Qubit™ dsDNA HS Assay Kit and was detectable in all samples but negative controls with an average of 1.3 ng/µl.

### Bacterial community analysis of the timeline survey by *16S rRNA* barcoding

#### Samples

We evaluated the bacterial phyllosphere diversity through Illumina sequencing of the *16S rRNA* ribosomal gene using primers 799F-1115R targeting the V5-V6 region and excluding chloroplastic DNA (9). *16S rRNA* gene amplicon sequencing was performed on 46 phyllosphere samples from 13 trees from both forests sampled 3-4 times throughout the 2018 growth season (**Dataset S1b**). We included one negative control (see section Sample collection) and one positive control (METH community) consisting in mixed genomic DNAs from 18 *Methylobacterium* isolates representative of diversity in SBL and MSH (**Dataset S1c,f**), one

*Escherichia coli* strain and one *Sphingomonas sp.* isolate from MSH (isolate DNA022; **Dataset S1c**).

#### Library preparation and sequencing

PCRs contained 1µL of sample DNA, 5 µL of Phusion Hot Start II Buffer (thermoscientific Fisher®), 0.5 µL of dNTP mix 10 µM, 0.5 µL of each primer at 10µM, 0.75µL of DMSO and 0.25 µL of Phusion Hot Start II Buffer DNA Polymerase (thermoscientific Fisher®) for a final volume of 25µL. PCRs were carried out in a thermocycler MasterCycler ProS Eppendorf© with the following steps: 30″ at 98 °C; then, 35 cycles of 15″ at 98 °C, 30″ at 64 °C and 30″ at 72 °C; and a final extension of 10′ at 72 °C. PCR products were loaded on 1% agarose gel ran at 120V for 45′ to control for amplification and fragment size with a 100 bp ladder (Invitrogen™). For each sample and the ladder, 1 µL of PCR product was mixed with 5 µL of dye buffer EZ-VISION® TREE (VWR™).

Samples were randomized before PCR amplification (random attribution of barcodes). PCR amplification (~400bp) was obtained for all samples and positive control but not negative controls. The *16S rRNA* amplicon library was prepared according to QIAseq FX DNA Library Kit (QIAGEN) protocole. The library was controlled for DNA concentration (Qbit™), quality (qPCR, NEBNext® Library Quant Kit for Illumina) and size distribution (Bioanalyzer DNA High Sensitivity, Agilent). Library concentration was adjusted according to the qPCR value at 6pM and sequenced with 1% phiX on Miseq (Illumina) with Miseq reagent kit v3 600-cycles (Paired-end 300 pb).

#### Amplicon Sequence Variant (ASV) definition with dada2

We obtained an average of 32,159 (range 12,162-55,181) paired reads per phyllosphere sample, 33,528 for the METH community (positive control) and 1,545 for the negative control. We processed *16S rRNA* reads in order to obtain an ASV (Amplicon Sequence Variants) abundance table per sample, using package *dada2* in R (10) with following modifications. According to sequence quality profiles, 3’ ends of forward and reverse reads from each sequence were trimmed 50 and 100bp, respectively (option truncLen in function *filterAndTrim*). Additionally, 5’ ends of both reads were trimmed 20bp in order to remove the primer part (option trimLeft in function *filterAndTrim*). After trimming, reads with expected error higher than 2 (option maxEE in function *filterAndTrim*) were discarded. Sample inference was performed using pseudo-pooling of samples (option pool=pseudo in function *dada*). Forward and reverse reads were merged together (function *mergePairs*) and chimeras were removed (function *removeBimeraDenovo*). After quality filtering (27% of sequences discarded), merging (13% discarded) and chimera removing (14% discarded), we conserved an average of 14,871 (range 8,096-21,529) sequences per phyllosphere sample, 16,367 for the METH community and 1,077 for the negative control, for a total of 24,733 unique variants (ASVs).

#### Contaminant ASVs filtering and rarefaction

We assigned taxonomy of each ASV identified in negative control and METH community using blast against NCBI nucleotide collection (nr/nt), uncultured/environmental samples excluded. In negative control (22 ASVs), 99.6% of diversity (proportion of sequences) corresponded to taxa typically associated with human oral and skin microbiomes (*Saccharibacteria*, *Leptotrichia, Corynebacterium*, *Actinomyces*, *Cutibacterium*, *Pseudoglutamicibacter*, *Prevotella* and *Kocuria*). In METH community (47 ASVs), 61.6% of diversity (proportion of sequences) corresponded to genomic DNA from bacterial orders mixed in the community: *Sphingomonadales* (1 ASV; 5.4% of diversity), *Enterobacterales* (*Escherichia coli*; 4 ASVs; 18.5% of diversity) and *Rhizobiales* (*Methylobacterium ssp.*; 18 ASVs; 36.4% of diversity). Although the remaining diversity was dominated by *Thermotogales* (*Fervidobacterium sp.*, 36.4% of sequences, 3 ASVs), these ASVs were absent from the negative control and phyllosphere samples, suggesting that these contaminants only affected the METH community. The remaining diversity corresponded to ASVs abundant in phyllosphere samples and thus likely corresponded to cross contamination among samples before the library preparation step. Each of these contaminant ASVs (*Thermotogales* excluded) had less than 0.3% of relative abundance in the METH community. We thus used this value as a conservative threshold to remove likely contaminant ASVs. In clear, any ASV that had a maximum relative abundance calculated across samples (negative control excluded) lower than 0.3% was discarded. We performed rarefaction curves on each sample to determine a conservative number of sequences to conserve per sample. Accordingly, the randomly picked 5,000 sequences per sample, hence excluding the negative control. After rarefaction, we obtained 732 ASVs, 725 of which were present in phyllosphere samples (42-292 per sample) and 24 in the METH community.

#### Taxonomic assignation of ASVs using SILVA

We aimed to assign taxonomy of each of the 732 identified ASVs (725 in phyllosphere samples), with emphasize on *Methylobacterium*, hence limiting taxonomic assignation at the genus level within *Methylobacteriaceae*, at the family level within *Rhizobiales*, at the order level within *Alphaproteobacteria*, at the class level within *Proteobacteria*, and at the phylum level within *Bacteria*. We used SILVA v.138 (11) as a database with *assignTaxonomy* function in R package *dada2* (**Dataset S1h**).

Among phyllosphere samples, 100% of diversity (proportion of sequences) was assigned to *Bacteria* (725 ASVs; **Dataset S1i**). The phyllosphere bacterial community was dominated in both forests by *Actinobacteria* (36.3% of sequences, 232ASVs), *Bacteroidota* (22.1%, 98 ASVs), *Deinococcota* (5.7%, 28 ASVs) and *Proteobacteria* (29.8%, 249 ASVs). *Proteobacteria* consisted in *Gammaproteobacteria* (9.1%, 87 ASVs) and *Alphaproteobacteria* (20.7%, 162 ASVs). In *Alphaproteobacteria*, more than 75% of diversity was found within the order *Rhizobiales* (13.1%, 72 ASVs), dominated by the family *Beijerinckiaceae* (13.0%; 67 ASVs). *Methylobacteriaceae* did not exist as a separate family in the SILVA database and *Methylobacterium* (annotated *Methylobacterium-Methylorubrum* (8)) was embedded within *Beijerinckiaceae* (15 ASVs, 1.3%). *Beijerinckiaceae* contained two “true” *Beijerinckiaceae* genera: *Methylocella* (4 ASVs, 0.5%), *Methylorosula* (3 ASVs, 1.5%), two incorrectly classified genera: *Psychroglaciecola* (*Methylobacteriaceae*; (12)), *Roseiarcus* (*Roseiarcaceae*; (13)) and one unknown genera annotated as *1174-901-12* but representing 9.5% of total diversity (38 ASVs). We blasted sequences from the two most abundant ASVs annotated “*1174-901-12* “ by SILVA against NCBI nucleotide collections (nr/nt, uncultured/environmental samples excluded) and obtained 100% identity with *Lichenibacterium minor* and *Lichenibacterium ramalinae,* respectively. These species were recently isolated from lichens colonizing birch trunks in boreal forests and represent the unique members of the newly described *Rhizobiales* family *Lichenibacteriaceae* (14). Finally, 5 ASVs (0.2% of diversity) were not assigned at the genus level.

*Methylobacterium* was present in almost all analyzed samples (45 out of 46), representing 1.3% [0.0-3.2%] of total sequence abundance. Using blast against NCBI databases (3) *refseq_genomes* and *refseq_rna* available for *Methylobacteriaceae* (Uncultured/environmental samples excluded), we determined that the 15 *Methylobacterium* ASVs identified by *16S rRNA* sequencing mostly belonged to clades typical of the phyllosphere: A9 (*M. phyllosphaerae/M. mesophilicum/M. phyllostachyos/ M. pseudosasicola/M. organophilum;* 0.87% of sequences, nine ASVs), A6 (0.29%; one ASV) and A1 (*M. gossipicola*; 0.13%, 3 ASVs; **Dataset S1i**). No ASV was assigned to group B or group C. We defined a new clade (A10) with two rare ASVs assigned to *M. komagatae* (<0.01%) but unrelated to any aforementioned clade.

##### *16S rRNA* community analyzes

We evaluated factors shaping phyllosphere microbial diversity estimated from *16S rRNA* barcoding, namely: forest of origin, host tree species and time of sampling. Relative abundances were normalized by Hellinger transformation to account for rare taxa, using function *decostand* in R package *vegan* (15). We performed a PERMANOVA to evaluate the contribution of each factor to microbial diversity (Hellinger transformation) using function *adonis* from R package *vegan* (**Table 1**; 10,000 permutations). As observed for PCA, forest of origin explained most of the variation (31.6%; $p<0.001$), followed by host tree species (15.6%; $p<0.001$) and sampling time (11.9%; $p<0.05$).

#### *Methylobacterium* isolation from a pilot survey in MSH in august 2017

##### Isolation

We performed *Methylobacterium* isolation from samples collected during the pilot survey in 2017 in MSH (57 samples from 19 trees; plots H0 and L0; 9-10 trees per plots; **Dataset S1b**). For each sample, we spread 10µL of phyllosphere microbial community glycerol stock (see section Sample collection) in a petri dish containing MMS synthetic solid media with 0.1% methanol sterilized by filtration (0.22µm filters), as sole carbon source to select for *Methylobacterium*, and 50mg/L of Cycloheximide to reduce fungal contamination (16). Petri dishes were incubated two weeks at 20 °C and 30 °C. Both temperatures were tested for all samples in order to minimize biases toward mesophylic isolates. From each petri dish, 0 to 3 pink colonies were isolated to maximize representativeness in term of color and size, spread separately on new MMS synthetic solid media (0.1% methanol, 50mg/L Cycloheximide) enriched with Sigma® RPMI1640 vitamins solution and 0.05g/L yeast extract to boost cell growth (17) and incubated 2-4 weeks at 30 °C or 20 °C according to the temperature of isolation. After incubation, colonies from petri dishes with contamination or with at least two different types of colonies (based on color) were spread on new MMS media whenever possible. Clean petri dishes were swabbed with 2mL of sterile distilled water. Colonies were collected in approximately 1mL of liquid and centrifuged in 1.5 mL Eppendorf tube, 10 minutes at 4°C (21130 rcf). Pictures of pellets were taken and tubes with evidence of contamination (two different colors in the pellet) were discarded. Clean pellets were resuspended; 450 µL were directly stored at -80 °C for future DNA extraction; 50µL were lysed by 10 minutes of incubation at 98°C, centrifuged 10 minutes at 4°C (21130 rcf) and stored at -20 °C for future isolate identification based upon amplification by PCR of marker genes, and 500µL were mixed with 500µL of glycerol 50% for culture stock (final glycerol concentration: 25%, storage at -80 °C).

##### *16S rRNA* V4 region amplification

We identified isolates from the pilot survey by PCR amplification and sequencing of the V4 region from *16S rRNA* ribosomal gene using primers 515F GTGCCAGCMGCCGCGGTAA (18) and 786R GGACTACHVGGGTWTCTAAT (19), universal for bacteria. For isolates DNA001-DNA024, identification was performed from genomic DNA extracted using DNeasy Blood & Tissue Kit (QIAGEN). For isolates LYS001-LYS096, identification was directly performed from cell lysate (see above). PCRs contained 1µL of cell lysates of genomic DNA, 5 µL of Phusion Hot Start II Buffer (thermoscientific Fisher®), 0.5 µL of dNTP mix 10 µM, 1 µL of each primer at 3µM, 0.75µL of DMSO and 0.25 µL of Phusion Hot Start II Buffer DNA Polymerase (thermoscientific Fisher®) for a final volume of 25µL. PCRs were carried out in a thermocycler MasterCycler ProS Eppendorf© with the following steps: 3′ at 98 °C; then, 35 cycles of 45″ at 98 °C, 1′ at 50 °C and 1′30″ at 72 °C; and a final extension of 10′ at 72 °C. PCR products were then sequenced by Sanger sequencing.

#### *Methylobacterium* isolate identification

We obtained *16S rRNA* nucleotide sequences from the V4 region for 80 pink colonies isolated for 18 out of 19 sampled trees (**Dataset S1c**). Sequences were manually curated according to the original chromatogram and classified using blast of the V4-*16S rRNA* sequence against NCBI databases (3) *refseq_genomes* and *refseq_rna* available for *Methylobacteriaceae* (Uncultured/environmental samples excluded). Four isolates did not closely match any *Methylobacteriaceae* reference and actually corresponded to *Stenotrophomonas sp*. (n=1), *Deinococcus sp*. (n=1) and *Sphingomonas sp.* (n=2). We assigned the 76 remaining isolates to previously identified *Methylobacterium* clades. We identified 8 unique V4-*16S rRNA* sequence variants. One variant present in 11 isolates was assigned to clade A9 (at least 99% of similarity with *M. phyllosphaerae/M. mesophilicum/M. phyllostachyos/ M. pseudosasicola/M. organophilum*); two variants in 19 isolates to clade A6 (*M. sp.*; 100% of similarity) and two variants in 12 isolates to clade A10 (*M. komagatae*; 100% of similarity). The three remaining variants identified in 34 isolates had 100% similarity with either clade A1 (*M. gossipicola*), A2 (*M. sp.*) or A3 (*M. sp.*). Because of the close V4-*16S rRNA* similarity (>99%) among some representative sequences of different clades, we affined the assignation of 24 representative isolates using sequencing and phylogenies of partial nucleotide sequences of two candidate marker genes: *sucA* and *rpoB* (Details in section Development of a *Methylobacterium*-specific molecular marker). We confirmed previous assignations to clades A6, A9 and A10 and distinguished isolates from clades A1 (n=30) and A2 (n=4; **Dataset S1c**).

#### Development of a *Methylobacterium*-specific molecular marker

##### Development of a fine-scale single-copy molecular marker specific to *Methylobacterium*

As an alternative to the *16S rRNA* gene, we developed a highly polymorphic marker targeting specifically – but not exclusively – isolates from the *Methylobacteriaceae* family. We choose two candidate genes, *rpoB* and *sucA*. Gene *sucA* is part of the 2-oxoglutarate dehydrogenase (OGDH) complex which catalyzes the decarboxylation of 2-oxoglutarate (20). It has been used as a marker gene to reconstruct phylogenies in *Salmonella* (21) and *Enterobacteriaceae* (22). Gene *rpoB* encodes the beta subunit of RNA polymerase and is widely used to reconstruct phylogeny in bacteria (1, 2, 22, 23). We retrieved complete *rpoB* and *sucA* nucleotide sequences available for

153 *Methylobacterium* genomes and 32 *Methylobacteriaceae* outgroups (*Microvirga*, *Enterovirga*; see **Dataset S1a**) and confirmed that both genes were single-copy in all genomes, contrary to *16S rRNA*. We performed alignment based on the amino-acid sequence in MEGA7 with default parameters (5). Based on the alignment, we identified five hypervariable (HV) regions (three in *sucA*, two in *rpoB*) flanked by well-conserved regions across *Methylobacteriaceae*, for which we designed specific primers (**Table S1**). We tested each primer pair targeting a HV region on 20 representative *Methylobacterium* isolates from the pilot survey. PCRs contained 1µL of cell lysates of genomic DNA, 5 µL of Phusion Hot Start II Buffer (thermoscientific Fisher®), 0.5 µL of dNTP mix 10 µM, 1 µL of each primer at 3µM, 0.75µL of DMSO and 0.25 µL of Phusion Hot Start II Buffer DNA Polymerase (thermoscientific Fisher®) for a final volume of 25µL. PCRs were carried out in a thermocycler MasterCycler ProS Eppendorf© with the following steps: 3′ at 98 °C; then, 35 cycles of 45″ at 98 °C, 30″ at 60 °C and 1′30″ at 72 °C; and a final extension of 10′ at 72 °C. PCR products were then sequenced by Sanger sequencing.

We successfully amplified and obtained clean sequenced for both *rpoB* HV regions in the 20 tested isolates (**Table S1; Dataset S1c,d**). We choose the first HV region targeted by primers Met02-352-F (AAGGACATCAAGGAGCAGGA) and Met02-1121-R (ACSCGGTAKATGTCGAACAG) as specific marker for *Methylobacteriaceae* for the rest of this study.

#### A consensus *Methylobacterium* phylogeny by coupling *sucA* and *rpoB* phylogenies

In order to validate *Methylobacterium* clade definition, we performed ML phylogenetic trees (100 permutations, complete deletion) for *rpoB* and *sucA* partial nucleotide sequences, separately. For *sucA*, we concatenated the three HV regions (1,663 bp) available for 189 reference genomes and the 14 tested isolates for which we obtained sequences for all of the three HV regions (**Figure S1a**). For *rpoB*, we concatenated the two HV regions (1,244 bp) available for 163 reference genomes and the 20 tested isolates (**Figure S1b**). We observed a strong congruence between both phylogeny topologies (summarized in **Figure S1c**). All defined clades were monophyletic in the *sucA* and *rpoB* phylogenies and supported by more than 80% of bootstraps in both for most clades but A1 (49 and 79%, respectively), B (99 and 68%, respectively) and A5 (89 and 45%, respectively). The consensus clade tree shows that group C is the more basal group of *Methylobacterium*. Among the remaining clades, we distinguished three groups of sister clades, among and within which phylogenetic relationships remain mostly unsolved: A1/A2/A3, A4/B and A5/(A6/((A7/A8)/(A9/A10))).

#### Culture-based assessment of *Methylobacterium* diversity of the timeline survey

##### Isolate isolation and identification

We monitored temporal and host-associated trends in *Methylobacterium* diversity in the phyllosphere by performing isolation on 28 trees sampled in 2018 in MSH and SBL (Timeline survey; **Dataset S1f**). Isolation was performed at 20 and 30 °C on MMS media with methanol as sole carbon source, as described in section Isolation**.** In a first batch, isolation was performed on all samples collected from 8 trees (4 per forest; 2 *Acer saccharum* and 2 *Fagus grandifolia*) between June and October 2018 (3-4 time replicates per tree). From this survey, we obtained 98 pink isolates (36 in MSH, 62 in SBL), for which we were able to amplify the *rpoB* marker using primers Met02-352-F and Met02-1121-R and get readable nucleotide sequences **(**details in section Development of a *Methylobacterium*-specific molecular marker). We obtained the highest average isolation success per sample in date 1 for MSH (27 June, n=5.0) and in date 2 for SBL (16 July, n=5.8). We thus selected these dates for a second isolation batch focusing on host-associated diversity. In each forest, we selected 10 trees sampled at the aforementioned dates and representative of diversity found in each forest. We repeated isolation and identification as described above and obtained 69 isolates (37 in MSH, 32 in SBL). Combining both batchs, we obtained 167 *Methylobacterium* isolates, 56.3% of which came from SBL, 43.7% from MSH; and 32.9% of which were isolated at 20 °C, 67.1% at 30 °C.

##### Isolate assignation to *Methylobacterium* clades

Among the 167 *Methylobacterium* isolates from the timeline survey, we identified 71 unique sequence variants, which is almost a 10-fold increase in comparison with V4-*16S rRNA* (see **section** *Methylobacterium* isolation from a pilot survey in MSH in august 2017). We assigned the 167 isolates to *Methylobacterium* clades (**Dataset S1f; Table S2)** using a phylogenetic tree (**Figure 3b**) inferred from an alignment combining partial nucleotide sequence for the *rpoB* marker sequenced for all isolates (first *rpoB* HV region), *rpoB* complete nucleotide sequences available for 185 *Methylobacteriaceae* complete genomes and partial nucleotide sequence for 2 *rpoB* HV regions we previously obtained for 20 representative *Methylobacterium* isolates from the pilot survey (1,244bp; detail in section Development of a *Methylobacterium*-specific molecular marker). Alignment was manually curated in MEGA7 (5), using the complete amino acid sequence as a guide, and sites not present in >70% of the sequences were removed using phyutility (v2.6). Nodal branch supports in the phylogenetic tree were estimated using MrBayes v. 3.2.7a as described in section Phylogenetics of plant-associated *Methylobacterium* diversity with the following modifications: standard deviation of split frequencies had stabilized to less than 0.05. Nodes with less than 30% of posterior probability were collapsed. the 167 isolates were assigned to clades within which their were embedded, using reference genomes and 20 isolates from the 2017 pilot survey as references. For these references, most clades were monophyletic, but clade A1 that formed a monophyletic group with A3, and clade A5 that splitted in two monophyletic groups (A5a, A5b). Isolates were assigned to A1 (n=9; 5.4%), A2 (n=3; 1.8%), A6 (n=41; 24.6%), A9 (n=100; 59.9%), A10 (n=6; 3.6%), A5b (n=1; 0.6%) and B (n=7; 4.2%).

##### Visual scaling of the *rpoB* phylogenetic tree according to pairwise nucleotide similarity.

We aimed to assess *Methylobacterium* isolate diversity a different depths within the *rpoB* phylogenies, with more emphases on the tips of the tree. We normalized the phylogenetic tree (**Figure 3b)** so it was scaled proportionally to nucleotide pairwise similarity (*PS*;). In other terms, we aimed to find a visual consensus between tree topology and evolutionary rate (here assumed to be proportional with PS). First, we imported the phylogenetic tree in Newick format in R using function *read.tree* in package *ape* (24). We converted the *read.file* object in a matrix filed with node names, with isolates in rows and nodes in columns. Nodes were ordered from the tip to the root of the tree, and then stacked to the root, so that the last column corresponded to the root of the tree, the before last column contained the closest embedded node(s), and so on. Each column of the matrix was labeled with a level number, $L$=1 corresponding to tips of the tree (sequences). For each node, we also calculated the median PS value ($PS$ = 1-*pdistance*) among embedded sequences, using a *pdistance* matrix calculated for all possible pairwise sequences in MEGA7 (complete deletion). *PS* values were transformed in inverse log scale ($PSc$= -log(1.005-$PS$)) to optimize resolution at the tips of the tree. Hence, each node was associated with a $L$ and a $PSc$ value. We used Pearson's correlation coefficient ($r^2$) between log($L$) and $PSc$ as an estimator of tree scaling ($r^2$ =-0.6071 for the initial tree). Then we iteratively moved nodes among levels in the matrix while respecting level hierarchy, until $r^2$ stabilized close to -1 ($r^2$=-0.9948; 3,000 iterations), meaning that *L* was roughly proportional to *PS*. We used the resulting matrix as guide tree for graphical representations in next sections.

#### PERMANOVA test at different depths in the *rpoB* phylogenetic tree

We aimed to test for association between *Methylobacterium* isolate diversity assessed at different depths within the *rpoB* phylogeny with sampling and isolation characteristics as proxy for *Methylobacterium* adaptive response to environmental variables through their evolution (**Figure 3a**). For each *PS* value in the phylogenetic tree in the range 0.950-1.000 (roughly corresponding to *PS* range within clades), we classified isolates into discrete taxa and performed a PERMANOVA (10,000 permutations) on *Methylobacterium* community dissimilarity using the *Bray-Curtis* index (*BC*) based on taxa absolute abundance (Hellinger transformation) using R package *vegan* (15). We tested for the relative contribution of four factors and their interactions on taxon frequency: forest of origin (*F*); temperature of isolation (*T*); sampling time (*D*) and host tree species (*H*). For each test, we performed permutations among isolates within batches of isolation to limit batch contribution in the explained variance (**Dataset S1j**).

#### Permutation test for node association

We asked specifically which nodes within the *Methylobacterium* phylogenetic tree were associated with the two major factors contributing to overall diversity, namely forest of origin and temperature of isolation (**Figure 3a**). For every level in the *rpoB* tree, we independently tested for association between taxa abundance (see above) and forest of origin (SBL and MSH) or temperature of isolation (20 and 30 °C) by permutation (100,000 permutations per level and per factor; **Figure 3b**). For each category of association (node-factor), we calculated p-values according to the following formula: $p=(b+1)/(m+1)$ where *b* was the number of expected values higher than the observed value and *m*, the number of permutations (25) and applied Bonferroni correction on p-values. To test for association with forest of origin, permutations were performed among isolates obtained at the same temperature and from the same batch of isolation. To test for association with temperature of isolation, permutations were performed among isolates from the same forest and from the same batch of isolation.

#### *Methylobacterium* community analysis of the timeline survey by *rpoB* barcoding

##### Samples, library preparation and sequencing

We evaluated the *Methylobacteriaceae* phyllosphere diversity through Illumina sequencing of the Methylobacteriaceae-specific marker specific using primers Met02-352-F and Met02-1121-R targeting the first hypervariable region of gene *rpoB* **(**details in section Development of a *Methylobacterium*-specific molecular marker). Amplicon sequencing was performed on 184 phyllosphere samples from 53 trees representative of diversity found in MSH (n=26) and SBL (n=27), and 48 of which allowed a monthly monitoring of diversity (3-4 samples per tree; **Dataset S1b,g**).

Library preparation and sequencing were performed as described in section Bacterial community analysis, with following modification. PCR amplification, library preparation and sequencing were proceeded in four different libraries, each containing a random subset of 46 phyllosphere samples, one negative control (see section Sample collection) and one positive controls (METH community; see section Bacterial community analysis). PCRs contained 1µL of sample DNA, 5 µL of Phusion Hot Start II Buffer (thermoscientific Fisher®), 0.5 µL of dNTP mix 10 µM, 0.5 µL of each primer at 10µM, 1.5µL of DMSO and 0.25 µL of Phusion Hot Start II Buffer DNA Polymerase (thermoscientific Fisher®) for a final volume of 20µL. PCRs were carried out in a thermocycler MasterCycler ProS Eppendorf© with the following steps: 30″ at 98 °C; then, 35 cycles of 15″ at 98 °C, 30″ at 60 °C and 60″ at 72 °C; and a final extension of 10′ at 72 °C. PCR amplification (~800bp) was obtained for all samples and positive controls but not negative controls.

##### Amplicon Sequence Variant (ASV) definition with dada2

We obtained an average of 30,423 (range 7,510–70,039) paired reads per phyllosphere sample (*n*=184), 29,297 (range 13,368–37,491) for METH communities (*n*=4) and 2,226 (range 467–3,490) for negative controls (*n*=4). We processed *rpoB* reads in order to obtain an ASV

(Amplicon Sequence Variants) abundance table per sample, using package *dada2* in R (10) with following modifications. Read trimming, learning error and concatenating steps were processed separately for each sequencing run (*n*=4). According to sequence quality profiles, 3' ends of forward and reverse reads from each sequence were trimmed 50 and 100bp, respectively (option truncLen in function *filterAndTrim*). Additionally, 5' ends of both reads were trimmed 20bp in order to remove the primer part (option trimLeft in function *filterAndTrim*). After trimming, reads with expected error higher than 2 (option maxEE in function *filterAndTrim*) were discarded. Sample inference was performed using pseudo-pooling of samples (option pool=pseudo in function *dada*). Because forward and reverse reads together (410 bp) did no cover the whole amplicon size (750 pb), they were concatenated together (function *mergePairs,* option *justConcatenate=T*), adding a "*nnnnnnnnnn*" string between forward and reverse sequences. Sequencing runs were combined (function *mergeSequenceTables*) and chimeras were removed (function *removeBimeraDenovo*). After quality filtering (29% of sequences discarded), concatenation (2% discarded) and chimera removing (27 % discarded), we conserved an average of 12,994 (range 3,935–25,807) sequences per phyllosphere sample, 13,915 (range 6,129–20,430) per METH community and 38 (range 7–100) per negative control, for a total of 44,518 unique variants (ASVs).

#### ASVs filtering and rarefaction

Before processing to rarefaction, we checked diversity within negative and positive controls (METH communities), using blast of the most abundant ASVs against NCBI RefSeq Genome Database (refseq_genomes, limited to *Alphaproteobacteria*) and Nucleotide collection (nr/nt), uncultured/environmental samples excluded. We recovered very few sequences in negative controls (7-100 sequences per replicate, 51 ASVs), most of ASVs present in only one replicate, suggesting very limited and scattered contamination. The most abundant ASV (45 sequences in a single replicate) corresponded to *Rhodococcus sp*., a typical contaminant of DNA extraction kits (26). Other ASVs (1-11 sequences per sample) were mostly assigned to *Rhizobiales* families typical of the phyllosphere (*Beijerinckiaceae*, *Lichenibacteriaceae*), suggesting very limited cross contamination between samples from a same sequencing run.

In METH communities (6,129-20,430 sequences per replicate, 243 ASVs), we found very good congruence in ASV absolute abundance across the four replicates (Pearson's correlation coefficient: 98.7-99.4%). Nucleotide sequences of the 19 most abundant ASVs (97.7% of total diversity) were exactly identical to *rpoB* partial sequences of 18 *Methylobacterium ssp.* (0.9-20.9% per ASV) and one *Sphingomonas sp.* (0.5%) isolates from which DNA was mixed to built the METH community. We thus considered them as "true ASVs". No ASV was assigned to *E. coli*, nor *Fervidobacterium sp.* although also present in the METH community and detectable through *16S rRNA* barcoding amplicon sequencing (see section Bacterial community analysis), confirming that the *rpoB* marker specificity is limited at least to *Alphaproteobacteria*. We considered that the 224 remaining ASVs (2.3% of diversity) could be either sequencing or PCR errors, contaminants or chimeric ASVs (i.e. resulting from concatenation of forward and reverse reads from different origins). We estimated the potential origin of these "false" ASVs by comparing their nucleotide sequences with those from the 19 trues ASVs, as well as homologous sequences from nine reference genomes that we identified through quick blast search of sequences from the most abundant false ASVs, and belonging to *Caulobacterales* (*n*=2) and *Rhizobiales* families *Beijerinckiaceae* (*n*=2), *Bradyrhizobiaceae* (*n*=2), *Lichenibacteriaceae* (*n*=2) and *Methylocystaceae* (*n*=1). For each putatively false ASV, we calculated nucleotide pairwise similarity (PS) between its nucleotide sequence and all references sequences. In order to identify chimeric ASVs, we did it separately for forward and reverse reads. We identified 12 relatively abundant chimeric ASVs (1.1% of total diversity) merely corresponding to scattered combinations of forward and reverse reads from true *Methylobacterium* ASVs. Other false ASVs corresponded to either *Methylobacterium* errors variants of true ASVs and/or contaminant (38 ASVs, 0.5% of total diversity), likely contaminant from phyllosphere samples (*Beijerinckiaceae*, *Bradyrhizobiaceae*, *Lichenibacteriaceae* and *Caulobacterales*; 77 ASVs, 0.5% of total diversity), chimeric combinations of ASVs among the aforementioned taxa (41 ASV, 0.1% of total diversity) and ASVs unrelated to any of the reference sequences (PS<90%; 56 ASVs, 0.2% of total diversity). Taken separately, false ASVs never exceeded 0.25% of relative abundance in one of the four replicated METH communities, while true ASVs always have at least 0.62% of relative abundance.

Accordingly, we filtered out ASVs that did not have at least 0.5% of relative abundance in a least one phyllosphere sample. We performed ASV filtering and rarefaction as described in section Bacterial community analysis, with a random sampling of 3,000 sequences in each phyllosphere sample and METH community (negative controls excluded; **Dataset S1k**). After rarefaction, we conserved 1,400 ASVs, with an average of 203 (range 48-355) per phyllosphere sample and 29 (range 25-35) per METH community. In METH communities, chimeric ASVs were correctly filtered out and remaining "false" ASVs (25 ASVs; 0.52% of total diversity) mostly corresponded to contamination from the most abundant taxa found across phyllosphere samples (see next section).

#### ASV taxonomic assignation

We assigned taxonomy of the 1,400 identified ASVs. We used a *rpoB* complete nucleotide sequence database available for *Bacteria* (44,673 reference sequences), previously developed by Ogier et al. (2) and last updated in December 2017. We formatted the database in R according to SILVA v.138 format (11) and processed to taxonomic assignation of the 100 most abundant ASVs with *assignTaxonomy* function in R package *dada2* (10). We considered assignation supported by at least 50% of bootstrap (*minBoot* =50). The 100 most abundant ASVs were assigned at the Class level to *Alphaproteobacteria*, and at the Order level to *Rhizobiales*, confirming specificity of the *rpoB* marker for this order. However, only 31 ASVs were assigned at the family level, to *Methylobacteriaceae* (28 ASVs) and *Bradyrhizobiaceae* (3 ASVs), respectively. We retrieved nucleotide sequences of the 13 most abundant ASVs that were not assigned at the family level and performed blast against NCBI databases (3) *refseq_genomes* and *refseq_rna* available for *Alphaproteobacteria* (Uncultured/environmental samples excluded). Nine ASVs matched *rpoB* sequences from at least one of five Rhizobiales genomes recently added in the NCBI databases (Query Cov > 99%; Per. Ident >98%): two from *Lichenibacterium* (*Lichenibacteriaceae*), a newly described genus (14), and three from *Group RH* a new, yet unnamed, *Beijerinckiaceae* genus (27). Accordingly, we reduced the *rpoB* database to *Alphaproteobacteria* to decrease computation time, and included the five aforementioned sequences. Additionally, we removed all sequences annotated as *Methylobacteriaceae* and replaced them by complete *rpoB* sequences from *Methylobacteriaceae* genomes available on September 2020 (**Figure 2**; **Dataset S1a**) combined with *rpoB* partial nucleotide sequences available from 20 isolates from the pilot survey (**Dataset S1c**). We annotated *Methylobacterium* reference sequences at the species level according to clades. The final *rpoB* database for *Alphaproteobacteria* contained 3062 reference sequences that we used to assign taxonomy of the 1,400 ASVs.

Most of ASVs (1,132) and diversity (94.91% of sequences) were assigned to *Rhizobiales* and 198 to *Caulobacterales* (4.32% of diversity). Within *Rhizobiales*, diversity was mostly assigned to *Beijerinckiaceae* (131 ASVs, 22.53% of diversity), *Lichenibacteriaceae* (262 ASVs, 25.68% of diversity) and *Methylobacteriaceae* (231 ASVs, 23.78% of diversity). A large proportion of *Rhizobiales* diversity (360 ASVs, 21.20% of diversity) was not assigned at the family level. To validate ASV taxonomy, we performed a phylogeny of 1,344 ASVs that could be exactly aligned upon they amino-acid sequence (*Rhodobacterales*, *Rickettsiales*, *Rhodospirillales*, some *Sphingomonadales* and unassigned *Alphaproteobacteria* ASVs excluded; **Figure S5a**). The ML tree (200 replicates) shows good support of main taxonomic groups identified with the *rpoB* database. Most of unassigned *Rhizobiales* diversity was found at the tip of long branches within monophyletic groups with few of no reference sequences: *Lichenibacteriaceae*, *Beijerinckiaceae* (*Group RH*) and *Methylobacteriaceae* (*Enterovirga*) clades, and in a monophyletic group sister of *Lichenibacteriaceae* and *Beijerinckiaceae*, and emcompassing some ASVs assigned to *Methylocystaceae* (*Methylocystaceae*-like group). We corrected ASV taxonomy according to their phylogeny (**Dataset S1k**).

After phylogenetic correction of taxonomy, ASV diversity was mostly found within *Caulobacterales* (209 ASVs, 4.42% of diversity) and within *Rhizobiales* (1,133 ASVs, 94,96% of diversity) in families *Methylobacteriaceae* (283 ASVs, 24.65% of diversity), *Beijerinckiaceae* (*GroupRH*; 165 ASVs; 24.58% of diversity), *Lichenibacteriaceae* (307 ASVs; 31.75% of diversity) and *Methylocystaceae-like* (171 ASVs, 11.04% of diversity). In *Methylobacteriaceae*, ASVs were mostly classified in *Methylobacterium* (200 ASVs, 23.05% of diversity), and *Enterovirga* (78 ASVs, 1.56% of diversity; **Dataset S1l**).

#### *Methylobacterium* ASV assignation to clades and of diversity between barcoding and isolation

We assigned the 200 *Methylobacterium* ASVs to clades using using a ML phylogenetic tree based on *rpoB* partial nucleotide sequences from the 283 Methylobacteriaceae ASV (including *Microvirga*: n=5 and *Enterovirga*: n=78), partial sequences including both *rpoB* variable regions available from 20 isolates from the pilot survey (**Dataset S1c,d**) and complete *rpoB* sequences from 185 references Methylobacteriaceae genomes (**Dataset S1a**). ASV Sequences were manually aligned to sequences from reference genomes based upon the amino-acid sequences. Missing positions in ASVs nucleotides sequences were replaced by "Ns". Both alignment and phylogeny were performed in MEGA7 (5). Only *Methylobacterium* ASVs embedded within nodes supported by at least 30% of bootstraps (200 permutations, pairwise deletion) and containing references assigned to a single clade, were assigned (**Figure S5b**).

We compared *Methylobacterium* diversity estimations from culture-dependant (*16s rRNA* and *rpoB* barcoding) and –independent methods (isolation). First, for *16s rRNA* barcoding (15 *Methylobacterium* ASVs) and *rpoB* barcoding comparison (200 *Methylobacterium* ASVs), we used data availalble for both methods from 41 phyllosphere samples from SBL (*n* = 27) and MSH (*n*=14) and the METH community (n=1; four replicates combined for *rpoB* barcoding). For each sample and each method, we calculated *Methylobacterium* ASVs sequence relative abundances (after excluding non-*Methylobacterium* ASVs) and combined these relative abundances for each clade (A1, A6, A9, A10 for *16S rRNA*; A1, A2, A3, A4, A6, A9, A10, B and unassigned ASVs for *rpoB*; number of ASVs per clade and method summarized in **Table S2**). We displayed results in a heatmap with samples in rows and clade relative abundance estimated by either *16S rRNA* or *rpoB* barcoding in columns using Euclidean distance to calculate similarity among relative abudances and samples **(Figure S5c**). Samples mostly clustered according to their origin (SBL or MSH). Relative abundances clustered according to clade (A1, A6, A9 and A10) rather than to method (*rpoB* or *16S rRNA* barcoding). We observed some inconsistency in clade A9 relative abundance when comparing both methods. This could be due to the high relative abundance of clade B detected by *rpoB* (19.1% of sequences) but not by *16S rRNA* barcoding (no sequence). By removing clades that were not detected by *16S rRNA* barcoding (B, A2, A3, A4, A5, unknown) in the calculation of clade relative abundance assessed by *rpoB* barcoding, we observed almost perfect correlation in relative abundance of clades A1, A6, A9 and A10 between both methods (data not showed). Second, for the culture-dependant (*rpoB* barcoding; 200 *Methylobacterium* ASVs from 184 phyllosphere samples) and –independent methods comparison (isolation; *rpoB* sequences from 167 isolates), we calculated pairwise nucleotide similarity (PS) among nucleotide sequences from isolates and ASVs by keeping only comparable regions (Ns removed; 333 bp left). We identified 123 isolates (out of 167: 73.7%) that had 100% identity (PS=1) with at least one ASV, for a total of 53 ASVs (out of 200), representing 71.2% of *Methylobacterium* diversity (sequence relative abundance) estimated from *rpoB* barcoding. Using a more relaxed threshold assuming up to 2 sequencing errors of mutations (PS>0.994), we found a match between 155 isolates (92.8%) and 124 ASVs representing 85.9% of diversity (**Figure S5d**). In both cases, we found a good congruence between relative abundances of sequence variants estimated from both methods (**Table S2**), indicating that our survey based on isolation was a good estimator of phyllosphere *Methylobacterium* diversity. The only exceptions were clade B, for which only 7 isolates (out of 167) were isolated in comparison with high relative abundance estimated from *rpoB* barcoding (19.1%), and clade A4 that represented 1.4% of *Methylobacterium* diversity based on barcoding but was not isolated. All analyses were conducted in R (28).

##### PERMANOVA analysis of *Methylobacterium* community dissimilarity

We performed PERMANOVA analyzes of 184 phyllosphere samples to evaluate relative contributions of forest of origin (*F*), plot within forest (*P*), host tree species (*H*), time of sampling (*T*) and their interactions, in *Methylobacterium* community dissimilarity (Bray distance). PERMANOVAs were conducted in R using function *adonis* from package *vegan* (15). *Methylobacterium* ASV absolute abundances (200 ASVs) were corrected by Hellinger transformation (*decostand* function) to account for rare ASVs and to correct for heterogeneity in *Methylobacterium* abundance between samples. For each analysis, 10,000 permutations were conducted between samples within randomized sequencing runs (*n*=4; *strata* option in *adonis* function) to control for variations due to sequencing errors. We tested following models: (i) a general model including all samples (*n*=184) and factors (*S**T*H*P*); (ii) two forest-specifics models conducted separately on MSH (*n*=85) and SBL (*n*=99) samples, hence excluding *S* from models (*T**H*P*; **Table 1**).

#### *Methylobacterium* ASVs association with environmental factors

We tested for the association of each *Methylobacterium* ASV with forest of origin (*F*), plot within forest (*P*), host tree species (*H*) and time of sampling (*T*) and their interactions (**Dataset S1m**). For each ASV independantly, we evaluated by ANOVA the contribution of these factors on the ASV relative abundance (*fx*; Hellinger transformation) under a linear model: lm($fx \sim S*T*H*P$). Then for each factor and interaction separately, we retrieve all *p*-values associated to the contribution (part of variance) to ASV relative abundance, and applied Bonferroni correction on *p*-values. In a principal component analysis (PCA) based on *Bray-Curtis* dissimilarity among 184 *Methylobacterium* communities (Hellinger transformation on 200 *Methylobacterium* ASV relative abundances), we reported ASVs contributions to the PCA only for ASVs significantly associated with one or either forest in the ANOVA (displayed in **Figure 4a**).

#### Spatial and temporal autocorrelation analysis on *Methylobacterium* ASVs

We quantified spatial and temporal dynamics of *Methylobacterium* community using autocorrelation analysis based on Bray-Curtis pairwise dissimilarity (*BC*) between 184 phyllosphere samples. In order to remove large-scale spatial variation due to strong difference in community composition between forests (**Table 1**), we analyzed MSH and SBL separately. For each possible pair of phyllosphere samples, we calculated *BC* on ASV relative abundance (200 ASV, Hellinger transformation), *pDist* as the spatial distance separating trees where the two samples came from (in meters) and *pTime* as the time separating dates when the two communities were sampled (in days). We evaluated the effects of *pDist* and *pTime* on *BC* under three different linear models by using ANOVA (**Table 2**). (i) *Spatial autocorrelation general models*: in MSH and SBL (two models), we evaluated the effect of *pDist* on *BC*. In order to take into account variations in community composition among dates, only pairwise comparisons among samples from a same date (*D*) were considered, and *D* as well as the *pDist:D* interaction were included in the model (**Figure 4b**). (ii) *Spatial autocorrelation models per date*. In each forest (n=2) and sampling date taken separately (n=4), we evaluated the effect of *pDist* on *BC* (eight models; **Figure 4e** only for MSH). (iii) *Temporal autocorrelation: general models*: in MSH and SBL (two models), we evaluated the effect of *pTime* on *BC*, regardless spatial scales (**Figure 4c**). For each model, we reported the average and standard deviation (*sd*) of the intercept, corresponding to the average and *sd BC* values among all the considered pairwise comparisons (BC distributions from model (ii) displayed in **Figure 4d**). For each factor (*pDist*, *Date*, *pDist:Date* and *pTime*), we also reported the average and standard deviation of estimates (slope), which significance was assessed by ANOVA on the linear model (**Table 2**).

#### *Methylobacterium growth performances* under four temperature treatments

We tested for the adaptive response of *Methylobacterium* isolates from the phyllosphere, to temperature variations during tree growing season. For 80 *Methylobacterium* isolated in 2018 in forests MSH (n=32) and SBL (n=47), we evaluated growth abilities under four temperature treatments, as a proxy for adaptation to temperature variations. Each treatment consisted in a first pre-conditioned (*P*) step during which each isolate was incubated for 20 days to either 20 (*P20*) or 30 °C (*P30*), and a second monitoring step (*M*) during which each pre-conditioned isolate was incubated and their growth monitored for 24 days at 20 °C (*P20M20* and *P20M30*) or 30 °C (*P30M20* and *P30M30;* **Figure S2**). We expected that treatments P20M20 and P30M30 mimicked stable thermal environments and that treatments P20M30 and P30M20 mimicked variable thermal environments.

#### Growing conditions

Pre-conditioning step (*P*; **Figure S2a,b**): For each isolate (*n*=80), two negative controls (*n*=2) and each temperature treatment (*n*=2), 10µL of cellular culture (thawed from -80 °C glycerol stocks) or sterile water for controls, were spread on 20mL of solid MMS media containing 0.1% methanol, 50mg/L of yeast extract and 50mg/L of vitamin mix (Sigma® RPMI1640). After 20 days of incubation at 20 °C (*P20*) or 30 °C (*P30*), petri dishes with no evidence of contamination were swabbed with 2mL of sterile distilled water. Colonies were collected in approximately 1mL of liquid and centrifuged in 1.5 mL Eppendorf tube, 10 minutes at 4°C (21130 rcf). Tubes with evidence of contamination (two different colors in the pellet) were discarded. Clean pellets were suspended by pipetting and cell concentrations were adjusted with sterile water to the same optic density $OD_{630}$=0.2, equivalent to about $1.6\times10^{8}$ cells/mL. No dilution was applied to negative controls.

Monitoring step (*M*; **Figure S2c,d**): For each pre-conditioned culture *P20* (*n*=82) and *P30* (*n*=82), 10µL (approximately $1.6\times10^6$ cells) were spotted on new petri dishes containing the same MMS media as used in the pre-conditioning step. Spots were distributed on petri dishes according to a 6×6 square grid (36 spots per plate), each dish containing 17 isolates and one negative control, for each of which one culture came from *P20* treatment and one from *P30*. *P20* and *P30* replicates from the same isolate were spotted next to each other in order to facilitate *P* treatment comparisons. Isolate positions were randomized elsewhere. Each petri dish was duplicated, one copy for incubation at 20 °C (*M20* treatment) and one for incubation at 30 °C (*M30*).

For each isolate and each combination of treatments (*PXXMXX*), we realized 5 replicates, randomly distributed in two series (*PXXM20* and *PXXM30*) of 24 petri dishes (**Figure S2e**). Within each *M* treatment, each petri dish had a different set and display of isolates, and replicates from the same isolate were in different petri dishes and in different positions. During the monitoring step, we took pictures of each petri dish with a Nexus LG device, at days 7, 13 and 24 after inoculation.

#### Image analysis

Pictures from each petri dish (*n*=24) and each time point (*n*=3) were first analyzed with ImageJ 1.52e software. Each original picture (**Figure S3a**) was converted in grey scale. Areas outside of the agar, as well as every visible particle other than bacteria spots within the agar area, were manually cropped using the elliptic tool. The picture was duplicated (**Figure S3b**). The first copy was used to measure raw bacteria spot intensities (*BW*). In the second copy, used for background correction (*BACK*), bacteria spots were cropped using the elliptic tool, by keeping as much background area as possible. Both copies having exactly the same dimensions (about 1,800x1,800 pixels per petri dish) were converted in matrix of pixel intensities using the */transform/image_to_results* tool and normalized in R in a 500x500-pixel matrix by averaging intensities. Cropped areas and pixels with intensities out of the range 50-200 were considered as missing values in *BACK* and *BW*. In order to reconstruct background intensities, missing values in the *BACK* file (including cropped positions of bacteria spots) were predicted from values with known intensity (**Figure S3c**). First, in order to tighten mesh in cropped areas, 15,000 pixels with missing intensities were randomly sampled and their intensity predicted from average known intensities in a 30x30 pixels windows. Second, intensities for remaining pixels with missing values were predicted in row (X-axis) and column (Y-axis), separately according to known and predicted values in 50-pixel windows, using function *runmean* (package *CaTools*). Third, for each pixel, predicted values in X and Y axis were averaged. Reconstructed intensities of the *BACK* files were then subtracted to the *BW* file in order to remove background (**Figure S3d**). For each spot, the growth area was determined as followed (**Figure S3e,f**). X and Y coordinates of the approximate central position of the bacteria spot was manually retrieved from the corrected *BW* file (36 per picture) loaded in ImageJ. In R, a 30x30-pixel window centered on these coordinates was defined. For each possible circle inscribed within this area, pixel intensity distribution outside and within the circle were compared using a t-test (minimum circle area: 200 pixels). The circle with the largest $t$ value was considered as the approximate profile of the bacteria spot, for which the average growth intensity (within the circle) and the average background intensity (outside the circle, within the 30x30-pixel window) were reported.

For of each spot, the average intensity was corrected by subtracting average local background intensity (**Figure S3f**). After correcting for local background, spot intensities from negative controls were almost indistinguishable from background ($I$ = 0.10±0.29).

We observed in average slightly but consistent higher spot intensity at the border ($I$= 5.8±5.5) than at the center of petri dishes ($I$ = 3.5±5.6), regardless of petri dish, isolate identity, time point, temperature treatment or other factors, suggesting that replicates located close to the border took advantage of less competition for nutrients (**Figure S3g**). Although positions of replicates from the same isolate were randomized, some could just by chance be systematically located close to the center or the border of a dish. We thus corrected for this border effect by predicting $I$ values in function of their position in the petri dish with the polynomial regression: $I{\sim}X^2Y^2+X^2Y+XY^2+X+Y$, were $X$ and $Y$ are the average coordinates of the spot on the petri dish (**Figure S3h**). Residuals from the polynomial regression were used as corrected $I$ values (**Figure S3i**).

#### Growth profile analysis

On average, spot intensity increased between days 7 ($I$ = 4.66±4.11) and 13 ($I$ = 5.32±4.91), followed by a average decrease in intensity at day 24 ($I$ = 4.22±4.14), illustrating that after reaching a maximal intensity (or yield), isolates eventually underwent starvation because of nutrient depletion in their immediate environment. For each isolate, each replicate and each treatment, we estimated yield (*Y*) and the growth rate (*r*; (29)). Because our survey was limited to three time points (7, 13 and 24 days), we estimated those values from predicted growth curves in the range 0-36 days, assuming that intensity was null at day 0 ($I_0$=0) and that growth curves followed a log normal distribution (**Figure S4a**; (30)). For each spot, we estimated the predicted growth curve, in the range 0-36 days after inoculation, from the best log normal model fitting assumed ($I_0$) and known intensity values ($I_7$, $I_{13}$ and $I_{24}$). We defined *Y* as the maximal intensity predicted from the log normal growth curve and *r* as the inverse of *log+lag* times necessary to reach Y (**Figure S4b**). We observe a good congruence between *Y* values predicted from log normal curves and maximum intensities observed from the three time points (**Figure S4c**), as well as a good congruence between predicted lag+log time and time at which the maximum intensity was actually observed (7, 13 or 24 days; **Figure S4d**). For a majority of replicates (87%), the predicted *Y* was reached before the 24$^{th}$ day of incubation (**Figure S4e**). From this point, we discarded the remaining replicates and considered 79 out of 80 isolates for which average *Y* and *r* could be predicted from at least one replicate (**Dataset S1n**).

In order to assess factors affecting *Methylobacterium* growth abilities under different temperature treatments, we constructed linear models (*lm* function in R) predicting *Y* and *r* (log transformations to meet normal distribution) in function of clade assignement of isolates (*C*), forest of origin (*F*), host tree species (*H*), time of sampling (*D*), temperature of isolation ($T_I$; at which each isolate was isolated), temperature of incubation during pre-conditioning ($T_P$) and monitoring ($T_M$) steps, and all possible interactions between factors ($\log(Y$ or $r) \sim H*S*D*C*T_P*T_M*T_I$) and evaluated the contribution of these factors in log(*r*) and log(*Y*) using an ANOVA (**Table 3; Dataset S1o**).

### REFERENCES

1. Vos M, Quince C, Pijl AS, Hollander M de, Kowalchuk GA. 2012. A Comparison of rpoB and 16S rRNA as Markers in Pyrosequencing Studies of Bacterial Diversity. PLOS ONE 7:e30600.
2. Ogier J-C, Pagès S, Galan M, Barret M, Gaudriault S. 2019. rpoB, a promising marker for analyzing the diversity of bacterial communities by amplicon sequencing. BMC Microbiol 19:171.
3. Boratyn GM, Camacho C, Cooper PS, Coulouris G, Fong A, Ma N, Madden TL, Matten WT, McGinnis SD, Merezhuk Y, Raytselis Y, Sayers EW, Tao T, Ye J, Zaretskaya I. 2013. BLAST: a more efficient report with usability improvements. Nucleic Acids Res 41:W29-33.
4. Tamura K, Nei M. 1993. Estimation of the number of nucleotide substitutions in the control region of mitochondrial DNA in humans and chimpanzees. Mol Biol Evol 10:512–526.
5. Kumar S, Stecher G, Tamura K. 2016. MEGA7: Molecular Evolutionary Genetics Analysis Version 7.0 for Bigger Datasets. Mol Biol Evol 33:1870–1874.
6. Ronquist F, Teslenko M, van der Mark P, Ayres DL, Darling A, Höhna S, Larget B, Liu L, Suchard MA, Huelsenbeck JP. 2012. MrBayes 3.2: Efficient Bayesian Phylogenetic Inference and Model Choice Across a Large Model Space. Systematic Biology 61:539–542.
7. Grafen A. 1989. The phylogenetic regression. Philos Trans R Soc Lond B Biol Sci 326:119–157.
8. Green PN, Ardley JK. 2018. Review of the genus Methylobacterium and closely related organisms: a proposal that some Methylobacterium species be reclassified into a new genus, Methylorubrum gen. nov. International Journal of Systematic and Evolutionary Microbiology 68:2727–2748.
9. Redford AJ, Bowers RM, Knight R, Linhart Y, Fierer N. 2010. The ecology of the phyllosphere: geographic and phylogenetic variability in the distribution of bacteria on tree leaves. Environmental Microbiology 12:2885–2893.
10. Callahan BJ, McMurdie PJ, Rosen MJ, Han AW, Johnson AJA, Holmes SP. 2016. DADA2: High resolution sample inference from Illumina amplicon data. Nat Methods 13:581–583.
11. Quast C, Pruesse E, Yilmaz P, Gerken J, Schweer T, Yarza P, Peplies J, Glöckner FO. 2013. The SILVA ribosomal RNA gene database project: improved data processing and web-

based tools. Nucleic Acids Res 41:D590–D596.
12. Qu Z, Jiang F, Chang X, Qiu X, Ren L, Fang C, Peng F. 2014. Psychroglaciecola arctica gen. nov., sp. nov., isolated from Arctic glacial foreland soil. International Journal of Systematic and Evolutionary Microbiology, 64:1817–1824.
13. Kulichevskaya IS, Danilova OV, Tereshina VM, Kevbrin VV, Dedysh SN. 2014. Descriptions of Roseiarcus fermentans gen. nov., sp. nov., a bacteriochlorophyll a-containing fermentative bacterium related phylogenetically to alphaproteobacterial methanotrophs, and of the family Roseiarcaceae fam. nov. Int J Syst Evol Microbiol 64:2558–2565.
14. Pankratov TA, Grouzdev DS, Patutina EO, Kolganova TV, Suzina NE, Berestovskaya JJ. 2020. Lichenibacterium ramalinae gen. nov, sp. nov., Lichenibacterium minor sp. nov., the first endophytic, beta-carotene producing bacterial representatives from lichen thalli and the proposal of the new family Lichenibacteriaceae within the order Rhizobiales. Antonie van Leeuwenhoek 113:477–489.
15. Dixon P. 2003. VEGAN, a package of R functions for community ecology. Journal of Vegetation Science 14:927–930.
16. Green PN. 2006. Methylobacterium, p. 257–265. *In* Dworkin, M, Falkow, S, Rosenberg, E, Schleifer, K-H, Stackebrandt, E (eds.), The Prokaryotes: Volume 5: Proteobacteria: Alpha and Beta Subclasses. Springer, New York, NY.
17. Tani A, Sahin N, Matsuyama Y, Enomoto T, Nishimura N, Yokota A, Kimbara K. 2012. High-Throughput Identification and Screening of Novel Methylobacterium Species Using Whole-Cell MALDI-TOF/MS Analysis. PLoS ONE 7:e40784.
18. Turner S, Pryer KM, Miao VP, Palmer JD. 1999. Investigating deep phylogenetic relationships among cyanobacteria and plastids by small subunit rRNA sequence analysis. The Journal of Eukaryotic Microbiology 46:327–338.
19. Caporaso JG, Lauber CL, Walters WA, Berg-Lyons D, Lozupone CA, Turnbaugh PJ, Fierer N, Knight R. 2011. Global patterns of 16S rRNA diversity at a depth of millions of sequences per sample. Proceedings of the National Academy of Sciences of the United States of America 108 Suppl 1:4516–4522.
20. Frank RAW, Price AJ, Northrop FD, Perham RN, Luisi BF. 2007. Crystal Structure of the E1 Component of the Escherichia coli 2-Oxoglutarate Dehydrogenase Multienzyme Complex. Journal of Molecular Biology 368:639–651.

21. Bell RL, González-Escalona N, Stones R, Brown EW. 2011. Phylogenetic evaluation of the 'Typhimurium' complex of Salmonella strains using a seven-gene multi-locus sequence analysis. Infection, Genetics and Evolution 11:83–91.

22. Ee R, Madhaiyan M, Ji L, Lim Y-L, Nor NM, Tee K-K, Chen J-W, Yin W-F. 2016. Chania multitudinisentens gen. nov., sp. nov., an N-acyl-homoserine-lactone-producing bacterium in the family Enterobacteriaceae isolated from landfill site soil. International Journal of Systematic and Evolutionary Microbiology, 66:2297–2304.

23. Küpfer M, Kuhnert P, Korczak BM, Peduzzi R, Demarta A. 2006. Genetic relationships of Aeromonas strains inferred from 16S rRNA, gyrB and rpoB gene sequences. International Journal of Systematic and Evolutionary Microbiology, 56:2743–2751.

24. Paradis E, Schliep K. 2019. ape 5.0: an environment for modern phylogenetics and evolutionary analyses in R. Bioinformatics 35:526–528.

25. Phipson B, Smyth GK. 2010. Permutation p-values should never be zero: calculating exact p-values when permutations are randomly drawn. Statistical Applications in Genetics and Molecular Biology 9.

26. Reagent and laboratory contamination can critically impact sequence-based microbiome analyses | BMC Biology | Full Text.

27. Wegner C-E, Gorniak L, Riedel S, Westermann M, Küsel K. 2019. Lanthanide-Dependent Methylotrophs of the Family *Beijerinckiaceae*: Physiological and Genomic Insights. Appl Environ Microbiol 86:e01830-19, /aem/86/1/AEM.01830-19.atom.

28. R-Developement-Core-Team. 2011. R: A language and environment for statistical computing. ISBN 3-900051-07-0. R Foundation for Statistical Computing. Vienna, Austria, 2013. url: http://www. R-project. org.

29. Lipson DA. 2015. The complex relationship between microbial growth rate and yield and its implications for ecosystem processes. Front Microbiol 6.

30. Zwietering MH, Jongenburger I, Rombouts FM, Riet K van 't. 1990. Modeling of the Bacterial Growth Curve. Appl Environ Microbiol 56:1875–1881.
